# Dynamic Reversal of IT-PFC Information Flow Orchestrates Visual Categorization Under Perceptual Uncertainty

**DOI:** 10.64898/2025.12.17.695044

**Authors:** Zahra Abouhadi, Hamid Karimi-Rouzbahani

## Abstract

Categorization relies on a dynamic interplay between sensory representation and cognitive control, yet the classical view posits a fixed, feed-forward information flow from the inferotemporal (IT) cortex to the prefrontal cortex (PFC). Whether this hierarchical directionality adapts to cognitive context, such as perceptual certainty, remains a fundamental question in systems neuroscience. We investigated this by recording intracranial neural activity in monkeys performing a delayed match-to-category task. We developed a novel connectivity framework, Model-Based Representational Connectivity Analysis (RCA), to simultaneously track the content, timing, and directionality of information flow between IT and PFC while controlling for common task-general representations. Our results revealed that while both areas rapidly encoded task-relevant information, the directionality of information flow was highly modulated by stimulus certainty. For high-certainty stimuli (far from the category boundary), we observed the classical feed-forward flow from IT to PFC. However, for low-certainty stimuli (near the category boundary), this hierarchy dynamically reversed, with a dominant, early feedback flow from PFC to IT preceding the feed-forward sweep. This feedback signal carried content-specific information related to the ambiguous stimuli, suggesting a top-down mechanism recruited to refine sensory representations. These findings challenge fixed-hierarchy models of visual processing, providing mechanistic evidence that the brain dynamically reconfigures the interactions between sensory and executive areas as a function of perceptual difficulty. We propose that the PFC initiates a top-down biasing signal to the IT cortex when sensory evidence is ambiguous, serving as an adaptive, context-driven control mechanism.

## Introduction

Categorization is a fundamental cognitive process that allows the brain to assign distinct perceptual experiences to shared functional classes, enabling organisms to generalize across variable sensory inputs and respond flexibly in novel situations (Barrett & Miller, 2026). It supports essential behaviors such as recognition, prediction, and decision-making, forming the foundation of higher-order cognition. Categorization is not limited to humans; it has been robustly demonstrated across a wide range of species, including monkeys (Vogels, 1999; Freedman et al., 2001), mice (Goltstein et al., 2021; Reinert et al., 2021), pigeons (Herrnstein & Loveland, 1964; Roberts & Mazmanian, 1988; Wasserman et al., 1988), and even insects (Avarguès-Weber et al., 2011; Wyttenbach et al., 1996). These studies suggest that many species can probably learn abstract category boundaries, generalize category membership to novel stimuli, and flexibly adapt categorical rules based on changing task demands. Together, these findings position categorization as one of the core computations of the brain, conserved across evolution and species which is broadly studied through behavioral, computational, and neurophysiological approaches.

Achieving categorization requires the brain to balance invariant neural representation with flexible sensory generalization to treat physically different stimuli as equivalent when they belong to the same conceptual group (Barrett & Miller, 2026; Riesenhuber & Poggio, 2000). To successfully classify a stimulus, the brain must ignore irrelevant perceptual variability while simultaneously maintaining an abstract category boundary or rule that guides the final decision. This transformation of complex sensory input into a goal-directed decision is orchestrated by the interaction between sensory representation areas, most notably the Inferotemporal (IT) cortex, and executive control centers, such as the Prefrontal Cortex (PFC) (Desimone, 1996; Kar & DiCarlo, 2021; Tomita et al., 1999). The information flow between IT and PFC is therefore fundamental to robust categorization and decision-making.

Decades of research on primates have established the complementary functional roles of these two regions. The IT cortex is recognized as the key hub for forming complex, view-invariant object representations (Tanaka, 1996; Vogels, 1999), which are necessary for fine-grained discrimination (Riesenhuber & Poggio, 1999). In contrast, the PFC is essential for maintaining the abstract rules of the task (Miller & Cohen, 2001; Wallis et al., 2001; White & Wise, 1999), resolving conflict (Duncan, 2001; Mansouri et al., 2009), and implementing strategic control during periods of ambiguity or delay (Miller & Cohen, 2001; Cole et al., 2016). While IT provides the sensory evidence (Desimone et al., 1984; Kiani et al., 2007; Vogels, 2022), PFC provides the cognitive context (Freedman et al., 2003; Meyers et al., 2008; Seger & Miller, 2010). Their constant interaction ensures that the appropriate sensory evidence is selected and maintained against the backdrop of the current behavioral goal (Tomita et al., 1999; Hebart et al., 2018; Kar & DiCarlo, 2021), a process that requires a delicate balance between feed-forward (FF) and feedback (FB) signals.

Anatomical and electrophysiological studies have established a generalized cortical hierarchy characterized by a dominant feed-forward directionality, propagating from IT to PFC, particularly during the rapid initial sweep of perception (Felleman & Van Essen, 1991). This FF flow is critical for the initial acquisition and mapping of sensory input onto category representations (DiCarlo et al., 2012; Rust & DiCarlo, 2010). However, categorization is rarely purely feed-forward. Feedback projections, typically originating in PFC and terminating in IT, are understood to play a modulatory role, supporting selective attention and memory maintenance (Moore & Armstrong, 2003). A central, unresolved question is whether this task-demanded feedback signal is merely a general modulator, or if the entire IT-PFC information hierarchy can dynamically re-organize and even reverse its dominant information flow direction when cognitive demands are high (Barrett & Miller, 2026).

The ambiguity of sensory evidence provides a crucial test case for the flexibility of the cortical hierarchy. When a stimulus is highly ambiguous, or falls close to a category boundary, the sensory evidence provided by IT is weak or contradictory. Successful categorization under these low-certainty conditions requires the system to rapidly leverage top-down signals to resolve the sensory conflict. Theoretical proposals, such as the Reverse Hierarchy Theory (RHT) (Ahissar & Hochstein, 2004), suggest that perception under difficult, uncertain conditions relies on slower, top-down guidance from executive areas. A powerful parallel and complementary framework is the Hierarchical Predictive Coding (HPC) theory (Friston, 2005), which views the brain as a machine constantly minimizing prediction error. In this view, sensory ambiguity results in a sustained, large prediction error signal being propagated up the hierarchy from IT. The necessary top-down control signal from the PFC would thus represent a revised, high-level prediction that is selectively weighted (or given higher “precision”) to suppress the bottom-up error signal in IT, thereby forcing convergence on a definitive category percept. Yet, a time-resolved and content-specific demonstration of whether the PFC actively initiates such reconfiguration of information flow—where the dominant direction becomes PFC -> IT—as a mechanism for uncertainty resolution is currently missing.

To address this gap, new analytical tools are required. Traditional functional connectivity methods often measure summarized statistical coupling between areas (Bar et al., 2006; Bastos & Schoffelen, 2016; Dijkstra et al., 2017; Gregoriou et al., 2009; Liu et al., 2025) ignoring interactions between neural populations (Basti et al., 2020) and are typically agnostic to the specific *information content* being exchanged (Anzellotti & Coutanche, 2018). Therefore, they cannot distinguish the flow of general attention/task set from the flow of stimulus-specific evidence necessary for resolving ambiguity. Here, we overcame this limitation by employing high temporal resolution electrophysiological recordings in conjunction with Model-Based Representational Connectivity Analysis (RCA) (Karimi-Rouzbahani et al., 2021, 2022) which is an extension of model-free RCA (Goddard et al., 2016). Model-based RCA allows us to precisely track the amount of cognitive content—such as information related to abstract category features or stimulus certainty—transferred from one area to another over time, even while strictly controlling for task-general representations via partial correlation.

We investigated the IT-PFC communication loop in monkeys performing a complex delayed match-to-category task that parametrically manipulated stimulus certainty (Roy et al., 2010). We first used Representational Similarity Analysis (RSA) to characterize the population coding, recurrence, and timing in both regions. Crucially, we then applied the Model-Based RCA to determine the directionality and content of the inter-areal communication. We show that while both areas rapidly encoded the relevant task information, the information flow direction was profoundly modulated by stimulus certainty. Our key finding is that the classical IT -> PFC feed-forward flow dominates under high certainty, but low certainty triggers a rapid, dynamic dominance of feedback flow (PFC -> IT) that precedes the canonical feed-forward sweep. These results challenge fixed-hierarchy models of categorization and provide a mechanistic framework detailing how the PFC actively and dynamically initiates the information flow to guide sensory processing when faced with perceptual ambiguity.

## Materials and methods

### Dataset

#### Subjects

We re-analyzed a previously collected rigorous dataset from monkeys where the individual-neuron analyses of PFC neurons have been published only (Roy et al., 2010). Here we included data from IT as well as PFC neurons of the same two Macaca mulatta monkeys (weighting 8-10 kg). The animals were handled in accordance with the National Institutes of Health guidelines and the Massachusetts Institute of Technology Committee on Animal Care.

#### Behavioral and intracranial recording methods

Full details of recording procedures were provided in the original study (Roy et al., 2010). Briefly, responses of 154 IT and 549 PFC neurons were recorded as the monkeys performed a delayed match-to-category task. PFC recording chambers were stereo-taxically placed over the principal sulcus and anterior arcuate sulcus. IT recordings were conducted using FHC 4-electrode microdrives, inserted through the same cannula and recorded from inferotemporal cortex. Neural activity from IT and PFC was recorded simultaneously in each session using independently movable electrode drives. Firing activity was recorded from well isolated neurons. Digitized waveforms were then saved for off-line sorting into individual neurons using principal component analysis (Offline Sorter, Plexon, Inc., Dallas, TX).

#### Stimuli and task

Two cat and two dog prototype models were generated in 3D and the 2D stimulus images were generated by morphing different percentages of the four prototypes (Figure 1A). Specifically, a three-dimensional morphing system (Shelton, 2000) was used to generate linear blends of two constituent prototypes. This resulted in a 2D space of stimulus with 14 morphs within each category. Dividing the stimulus space into two different category schemes yielded Schemes A and B, where the boundary lines were orthogonal across the two schemes. In each categorization scheme, only two explicit categories were taught to the monkeys. A morphed stimulus was assigned a category if it contained more than 50% contribution from the prototype in that category.

**Figure 1.**
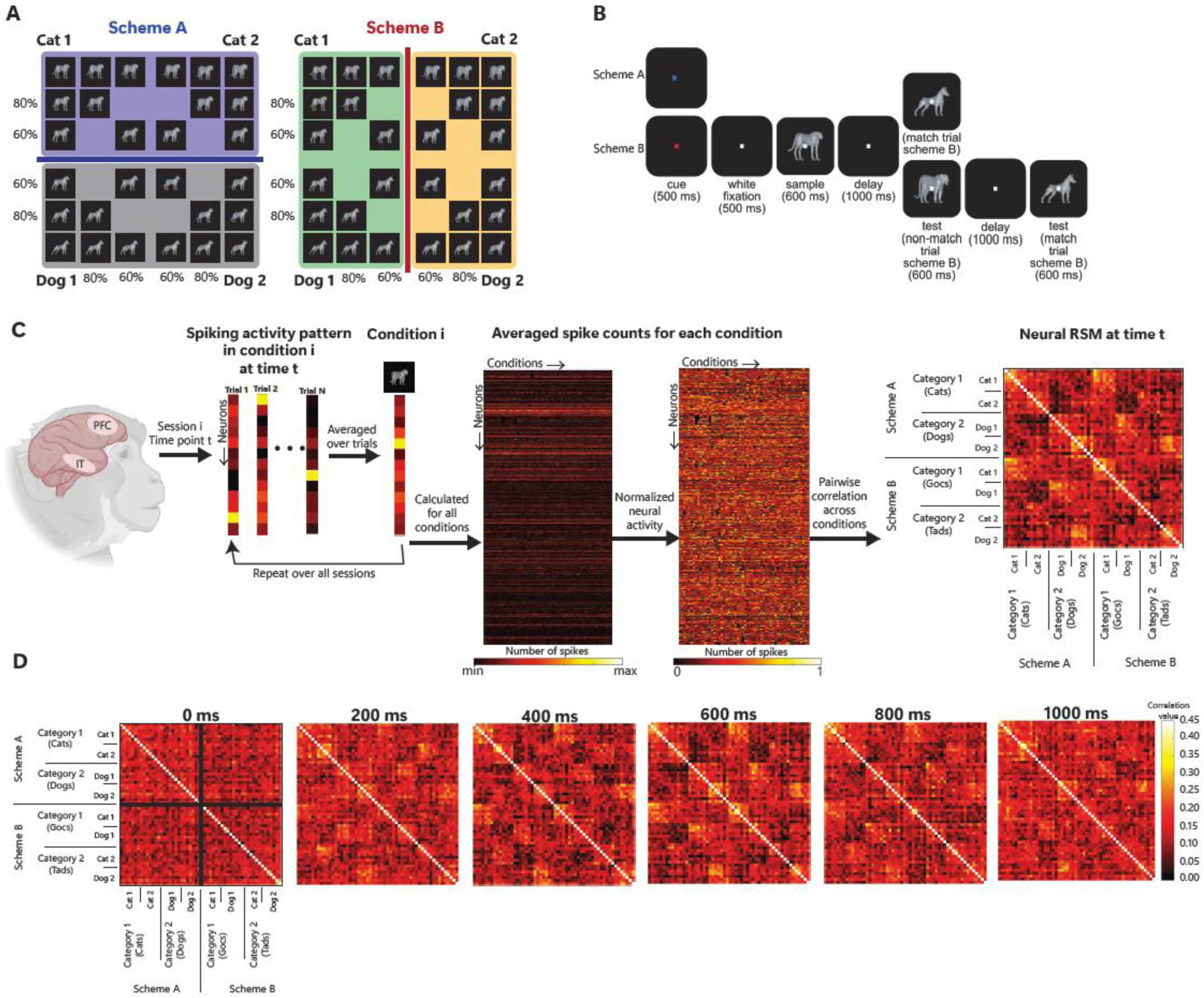
Stimuli, tasks and population-level representational similarity analysis (RSA). **(A)** Stimuli were created using a morphing system that linearly interpolated between four prototype images shown on corners of the stimulus matrix (stimuli re-generated upon provision by Roy et al., 2010. **(B)** Monkeys categorized stimuli using two orthogonal category schemes as shown by blue and red lines in A using a delayed match-to-category task. Trials began with a fixation, followed by a color-cued scheme. Monkeys viewed a sample stimulus, then judged whether a test image matched the category of the sample image (lever release for matches and hold for non-matches) after a delay. **(C)** Representational similarity analysis of population-level neural activity. Upon within-condition trial averaging and normalization of activities across neurons, a Spearman’s correlation was calculated between population-level neural activity patterns across conditions to form a representational similarity matrix (RSM) for every 100ms (5ms steps) time window over the trial. **(D)** Example neural RSMs at specific time points over the sample presentation window (averaged over the two monkeys), which go from a noisy pattern at time 0 to a more structured pattern later on the trial.

During training, monkeys viewed thousands of images based on a combination of the four prototypes. In recording sessions, 28 images each combining one of six ratios of prototypes were presented (ratios of 100:0, 80:20, 60:40, 40:60, 20:80, and 0:100). The images of the six morphing ratios and category assignments are shown in Figure 1A; in Scheme A, the two categories were named Cats and Dogs as they were generated on the line connecting cat and dog prototypes, while in Scheme B, they were named Gocs (left) and Tads (right) as they were orthogonal to the Cats and Dogs categories being influenced by cate and dog prototypes.

A delayed match-to-category task was performed by monkeys (Figure 1B). The task began with monkeys holding a lever and maintaining a fixation target for 1000 ms. The color of the fixation dot indicated which category scheme would be in effect for the trial (blue for Scheme A and red for Scheme B). At 500ms prior to the onset of sample presentation, the fixation reverted to white. Sample image was then presented for 600 ms, followed by a 1000 ms memory delay. After the memory delay, a test image was presented. A trial was labeled a *match* if the test image’s category matched the category of the sample, and a *non-match* if it did not. In a match trial, monkeys released the lever to receive a juice reward, and in non-match trials they continued to hold the bar through a second 1000 ms delay which was followed by a category-match image requiring the monkey to release the bar for a juice reward. Category scheme A/B and match/non-match trials were interleaved randomly and occurred at similar probabilities.

Scheme A was the first category scheme both animals were trained on which attaining a performance of 80% took ∼6 months. Then the monkeys were trained on category scheme B exclusively for around 4 months until the same performance criterion was achieved. Only then sessions including both schemes were introduced. Initially, blocks of 20 trials per scheme were presented which were reduced to one after 4 subsequent weeks enabling random selection of schemes.

### Data Analysis

All analyses were performed in MATLAB using custom codes (version 2020a).

#### Representational Similarity Analysis (RSA)

To evaluate the population-level neuronal activity in the brain, we adopted a Representational Similarity Analysis approach (Kriegeskorte et al., 2008). Specifically, we computed representational similarity matrices (RSMs) on every time point. These RSMs were constructed using activity patterns from all recorded neurons over 100 ms sliding windows with a sliding step of 5 ms across the time. Spike counts per time window were averaged over trials for each condition, yielding a [neurons × conditions] data matrix for generating RSMs. To avoid correlations from being driven by the absolute level of activity in highly active and/or inactive neurons, each neuron’s activity was z-scored across conditions prior to generating the RSMs, ensuring comparability of response magnitudes across the neural population. Then, we calculated the similarity between conditions (using Spearman’s correlation) and arranged them into an RSM. With 28 stimuli per category across two schemes, the RSM dimensions were 56 × 56 (Figure 1C).

#### Conceptual models

To evaluate how distinct types of information were represented in the neural activity patterns as summarized in the neural RSMs, we constructed ten conceptual model RSMs. Each of these models looked at the data from a different lens/persepective quantifying the amount of information about a specific aspect of the experimental condition in neural activity patterns (Kriegeskorte et al., 2008; Nili et al., 2014). Here, information is defined as *distinction between different experimental conditions (e.g., between different stimuli or between different schemes)*. In the models, similarity and dissimilarity between experimental conditions were represented by high and low values, respectively. The experimental conditions which were hypothesized to be more similarly encoded (e.g., *stimuli within the cat category*) were represented by the high value of 1 whereas the experimental conditions which were hypothesized to be less similarly encoded (e.g., *stimuli across cat vs. dog categories*) were represented by the low value of 0. We implemented two types of models: binary models, which included only zeros and ones assuming maximum distinction between conditions and non-binary models which had values between zero and one allowing for capturing intermediate distinctions between conditions. Below we summarize the models (Figure 2A):

**Figure 2.**
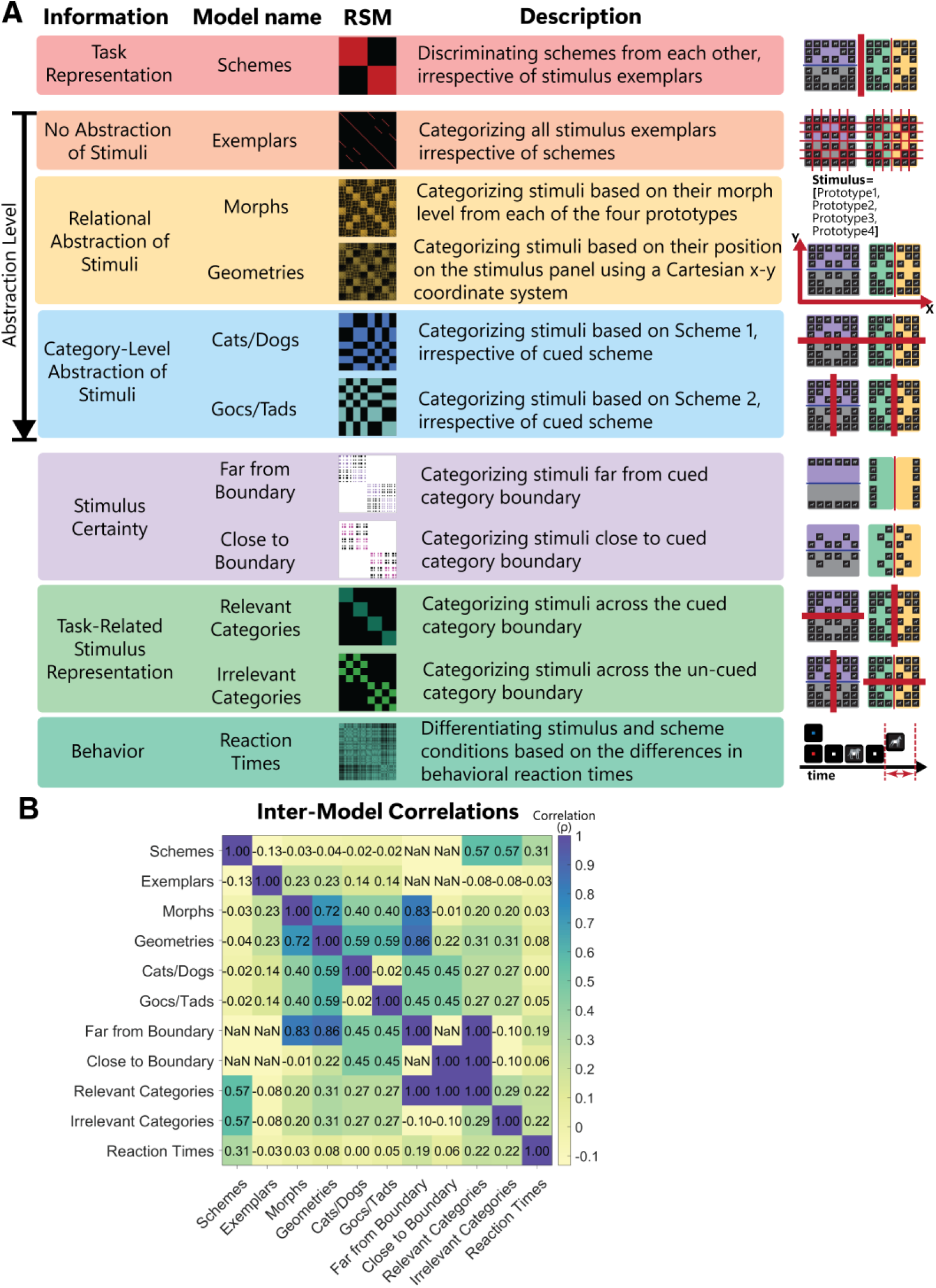
Conceptual models and their correlations. **(A)** Summary of the representational similarity models designed to quantify different types of information in neural population activity. The Information column specifies the information captured by each model (or pair of models). The Model name column provides model names used in the text. Each model’s similarity values are color-coded distinctly where black indicates 0 (least similar), colored elements show 1 (most similar), intermediate colors represent intermediate similarities, and white indicates condition pairs which were excluded from the analyses. The Description column details the different information each model quantifies. Finally, the right-most column illustrates the distinction that each model quantifies on the stimulus-morph table. **(B)** Spearman’s correlation between models.

##### Schemes Model

quantified scheme-related information by distinguishing the two schemes irrespective of stimuli. This model would reveal if the brain distinguished the schemes.

##### Exemplars Model

quantified exemplar-level stimulus information by distinguishing every single stimulus exemplar from the rest, irrespective of schemes. This model would reveal if the brain distinguished every single stimulus from the rest.

##### Cats/Dogs Model

quantified the discrimination of cats/dogs stimuli irrespective of the cued scheme. Note that monkeys were cued to make this categorization only in scheme A but not B. This model would reveal if the brain distinguished cats/dogs stimuli irrespective of the current cued scheme.

##### Gocs/Tads Model

quantified the discrimination of gocs/tads stimuli irrespective of the cued scheme. Note that monkeys were cued to make this categorization only in scheme B but not A. This model would reveal if the brain distinguished gocs/tads stimuli irrespective of the current cued scheme.

##### Morphs Model

which was a non-binary model, quantified the amount of information about morph levels across stimulus exemplars, irrespective of schemes. This model quantified the similarity between the morph levels which generated each stimulus exemplar. These morph levels were concatenated into a 4-element vector for each stimulus (reflecting the morph level from each prototype e.g., [1.0 0.0 0.0 0.0] for cat 1 stimulus, which is purely from cat 1 and [0.8 0.2 0.0 0.0] for the next stimulus to the right which is constructed by morphing 80% of cat 1 prototype and 20% of cat 2 and no contribution from the other two prototypes and so on) and then correlated across stimuli. Finally, all pair-wise correlations were normalized between 0 and 1. This model would reveal if the brain distinguished the stimuli based on the morph levels generating them.

##### Geometries Model

which was another non-binary model, quantified the similarity between stimulus pairs through measuring Euclidean distance in a 2D Cartesian space as reflected in the stimulus morph panel shown in Figure 1A. Accordingly, the stimuli which were horizontally side by side had a Euclidean distance of 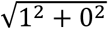 and cat 1 and dog 2 prototypes had a Euclidean distance of 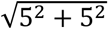 because they were 5 horizontal and 5 vertical steps apart on the stimulus morph panel. The final model was generated by computing correlations between inter-stimulus distance vectors, followed by normalizing the correlation values between 0 and 1. This model would reveal if the brain held a geometrical representation of stimuli which matched the stimulus morph table.

##### Abstraction

The above stimulus-based models capture different levels of stimulus abstraction. In concrete-to-abstract levels of abstraction, the Exemplars model was the most concrete as it considered any distinct pairs of stimuli as different irrespective of their categorical associations defined by the categorical schemes. This would define a very strict representation ignoring any relationship between stimuli. On the other hand, Cats/Dogs and Gocs/Tads models were abstract representations where they grouped together the stimuli based on the categorical schemes defined in the experiment. Note that, in these models, stimuli were not grouped based on current trial’s cued scheme but rather the two learned schemes irrespective of if they were cued or not on the current trial. Finally, to precisely characterize the coding of relational information, we included ’Semi-Abstract’ models (Morphs and Geometries) which capture the intermediate coding required for generalization, bridging the gap between concrete visual features and the abstract decision rule. Morphs and Geometries maintained the stimulus-level information by comparing inter-stimulus morph levels/distances while also discriminating schemes as stimuli which were more distinct/distant generally belonged to different categories (see the positive correlations between the Morphs and Geometries models and the other stimulus models in Figure 2B). This graded organization were evaluated based on the hierarchical transformation of visual into semantic information observed from lower to higher sensory areas including along the ventral visual hierarchy (DiCarlo et al., 2012).

##### Far from Boundary Model

quantified information about the stimuli far from category boundary in each scheme by discriminating stimuli at 100% morph level across the boundary, excluding other morph levels in analysis. This model would reveal if the brain representations distinguished stimuli far from the category boundary more strongly, supporting higher certainty.

##### Close to Boundary Model

quantified information about the stimuli close to category boundary in each scheme by discriminating stimuli at 60% and 80% morph levels across the boundary, excluding other morph levels. This model would reveal if the brain representations distinguished stimuli close to the category boundary less strongly, supporting lower certainty.

##### Relevant Categories Model

quantified information about cued stimulus categories across the category boundary according to the cued scheme. This means distinguishing cats vs. dogs in scheme A and gocs vs. tads in scheme B. This model would reveal if the categorical representations were enhanced when aligned with the cued scheme.

##### Irrelevant Categories Model

quantified information about un-cued stimulus categories across the category boundary according to the un-cued scheme. This means distinguishing cats vs. dogs in scheme B and gocs vs. tads in scheme A. This model would reveal if the categorical representations were deteriorated when orthogonal to the cued scheme.

##### Reaction Times (RT) Model

quantified how behavioral responses differed across stimulus and task conditions. Reaction times were calculated as the time delay between test stimulus onset and the monkey’s correct behavioral response. To construct the model, reaction times were subtracted across pairs of conditions. Finally, all values were normalized in the [0,1] range, enabling comparison with neural RSMs. This model would reveal if the differences in reaction times across conditions were reflected in the differences between neural representations of the same conditions. It is of note that this model is derived from behavioral data rather than any external hypothesis. Only match trials were included in the development of the RT model to avoid monkeys’ predictions affecting representations - after a non-match test image there always was a match image.

To quantify scheme information, we analyzed neural data aligned to the cue presentation, focusing on the period when the scheme was displayed (-200 to 1200 ms). On the other hand, to investigate the encoding of visual stimuli, data aligned to the sample onset was used. These analyses targeted both the initial stimulus processing period (-200 to 600 ms post-sample) and subsequent memory delay period (600-1600 ms). Models fit to these epochs included all models except Schemes and Reaction Time models. Finally, to link neural dynamics directly to the decision output, we quantified behavioral responses information using data aligned to the behavioral response onset.

Figure 2A summarizes the information that each model quantifies. While some of the hypothesized and designed models inevitably showed some correlations (Figure 2B), what is also critical is the non-overlapping representations and dynamics that each model captures in neural representations. Specifically, to address the inherent correlations between stimulus-driven features (e.g., morph levels, stimulus geometry) and task-driven variables (e.g., category labels, task schemes), we utilized a model-comparison and partial-correlation approach. Specifically, for all connectivity analyses (RCA), we calculated partial directed correlations to isolate the unique variance contributed by the source area to the destination area. Where task-set information could potentially confound stimulus-related results, the ‘Scheme’ RDM was partialled out of the analysis. Furthermore, the task’s dual-scheme design (interleaving Scheme A and B) allowed us to functionally dissociate sensory-driven representations from category-driven representations, as the categorical identity of identical stimuli shifted depending on the active task rule. This approach ensures that the reported informational flows reflect specific representational currencies rather than shared global variance.

#### Information Representations (encoding)

To quantify if neural activity patterns in IT and PFC aligned with our hypothesized models, we performed the following analyses. First, for both neural RSMs and conceptual models, we extracted the upper triangular elements (excluding the main diagonal) and vectorized them to create representational similarity vectors (RSVs). Next, at each time point, we computed *Pearson* correlations between neural RSVs and each model RSV. These correlation values quantified how strongly each information type was encoded in the neural data.

When correlating Relevant and Irrelevant category models with neural RSVs, we partialled out the Schemes model (using MATLAB partialcorr function) as it correlated with those models, to provide stimulus information under different schemes without being affected by the potential differences between schemes (see the high correlations between relevant/irrelevant models and the scheme model in Figure 2B).

#### Recurrence of Information Representations

In addition to information coding, we also developed an RSA-based method to quantify the recurrence of information within each area separately. This allowed us to determine how sustained information was within each region. Unlike traditional recurrence measures which were based on machine learning classification (King & Dehaene, 2014) and only evaluated binary contrasts, our novel approach is based on the same RSVs and the above-mentioned models to distinguish between recurrence of general information and the recurrence of information specific to the defined conceptual models. This was achieved by removing components of current neural representations that could be explained by prior representations within the same region, using conceptual models as lenses of information. This procedure, detailed in Equation 1, was performed independently for IT and PFC:

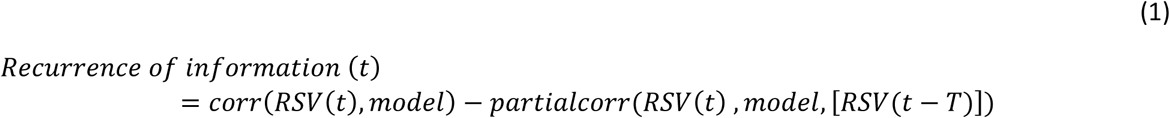

here, the variable in the bracket represents the partialled out RSV. 𝑇 denotes the temporal integration moment in the past, computing recurrence from onset to the current time 𝑡. As in the above, in the case of Relevant and Irrelevant category models, Scheme model was partialled out from both terms in equation (1).

#### Model-Based Representational Connectivity Analysis: Tracking Content-Specific Information Flow

Traditional functional connectivity analyses often measure statistical coupling (e.g., correlation, Granger Causality) between neural activity patterns, but they are agnostic to the *information content* being exchanged. To overcome this limitation, we developed *Model-Based RCA* (Karimi-Rouzbahani et al., 2021), which combines the power of Representational Similarity Analysis (RSA) with multivariate connectivity models. This approach specifically measures the amount of *information content* (e.g., ’Far’ certainty information or ’Close’ certainty information) transferred from region A to region B over time, ensuring our measure reflects the flow of cognitive variables rather than raw neural variance.

To measure directional flow of representations between areas, we first vectorized the upper triangular elements of each RSM (excluding diagonal elements) to create RSVs. We then quantified how much the representations in the source area in the past contributed to the representations in the destination at present time (measured by its correlation to the hypothesized model), while controlling for the past representations in the destination area (based on Granger causality principles). We quantified feed-forward (IT -> PFC) and feedback (PFC -> IT) flows using equations 2 and 3, respectively:

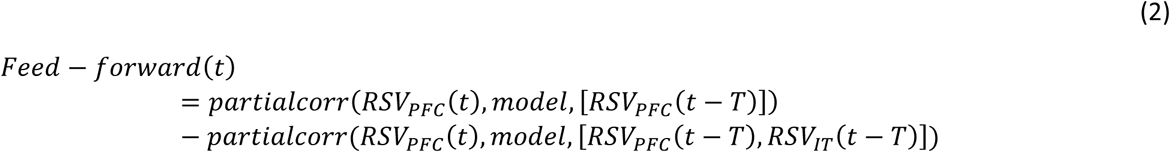

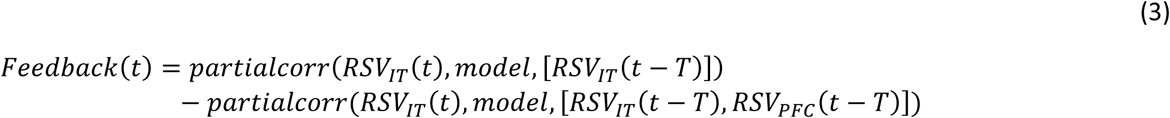

In equation (2) and (3), the delay time 𝑇 was considered to range from 5 to 150 ms to cover the most plausible flows between the two areas without capturing too much recurrence of the same information. For direct comparisons (e.g., Far vs. Close models) we averaged flow values over all delays yielding a curve representing average flow of information.

As an improvement to our original RCA method (Karimi-Rouzbahani et al., 2021), in the new RCA analyses, the past representations in the destination area were also partialled out to make the flow more specific to be driven by the source area. For the Relevant and Irrelevant Category models, the Scheme model was again partialled out to exclude scheme-dependent effects from information flow computations. These critical steps ensure that the measured flow reflects the transfer of stimulus-specific sensory evidence rather than merely the generalized attentional set or task rule being maintained in the PFC.

### Statistical Analysis

Statistical significance for the most analyses in this study was determined using non-parametric random permutation tests to generate empirical null distributions. Statistical tests were performed in MATLAB, and a significance threshold of p < 0.05 (cluster-corrected) was used.

#### Information Coding

To assess the statistical significance of information coding while controlling for the multiple comparisons problem inherent in time-resolved analyses, we employed a non-parametric permutation approach. For each time point, a null distribution was generated by shuffling condition labels 1000 times and recalculating the Spearman correlation between the neural Representational Similarity Matrices (RSMs) and the conceptual model. Time points in the original data were considered candidates for significance if the observed correlation exceeded the 95th percentile of the permutation-based null distribution (p < 0.05, one-tailed). To control the Family-Wise Error Rate (FWER) and account for the temporal dependency of neural signals, we applied a cluster-extent threshold. Only clusters of significant time points exceeding 100ms (20 consecutive 5ms bins) were reported. This duration was chosen based on the typical temporal stability of neural population codes and serves to effectively filter spurious, transient noise while preserving sustained representational states.

#### Recurrence of Representations

The statistical significance of representational recurrence was evaluated using a non-parametric permutation approach. At each time point, condition labels within the neural RSMs were permuted 1,000 times to generate a null distribution for the recurrence metric (Equation 1). Observed recurrence values were considered statistically significant if they exceeded the 95th percentile of the permuted distribution (p < 0.05, one-tailed). To identify highly persistent representational states and control for the family-wise error rate (FWER) inherent in cross-temporal analyses, we applied a stringent cluster-extent threshold within the temporal generalization matrix. Given the high dimensionality of the time-by-time generalization space, we required a minimum cluster mass of 2,000 contiguous significant bins (p < 0.05) within the analysis triangle for a result to be considered statistically significant. This conservative threshold ensures that reported effects reflect stable, long-term neural recurrence rather than transient, spurious correlations.

#### Representational Connectivity Analysis

The statistical significance of feed-forward and feedback information flows was assessed using a non-parametric permutation approach. For each time point and delay, condition labels within the neural RSMs were shuffled 1,000 times to generate a null distribution of the information flow metric (Equations 2 and 3). Observed flow values were considered significant if they exceeded the 95th percentile of their respective null distributions (p < 0.05, one-tailed). For the aggregate curves representing information flow averaged across delays (5–150 ms; 10 delays), we implemented a consistency-based significance criterion: a time point was deemed significant only if the underlying flow values reached significance (p < 0.05) in more than 50% of the individual delays contributing to that average. To control for the family-wise error rate across the time-delay space, we applied a dual-threshold cluster correction. Only clusters that spanned more than 20 consecutive time points and encompassed a cumulative total of 200 significant flow instances across the time-delay matrix were considered statistically robust.

We also used Bayes Factor (BF) analysis to determine the difference between flows across different information types. Specifically, we interpreted the levels of BF evidence more strictly than conventional thresholds (Dienes, 2014; M. D. Lee & Wagenmakers, 2005): BFs above 6 was considered evidence for the alternative (i.e., difference; H1) and BFs below 6 was considered insufficient evidence for the alternative hypothesis (i.e., no difference; H0). We used BF t tests to compare the level of flow for different types of information. Priors for all BF analyses were determined based on Jeffrey-Zellner-Siow priors (Jeffreys, 1998; Zellner & Siow, 1980), which were derived from the Cauchy distribution based on the effect size initially calculated in the algorithm using t tests (Rouder et al., 2012).

## Results

Monkeys successfully performed the delayed match-to-category task with high accuracy (Mean = 0.92; SD = 0.06) and fast reaction times (Mean = 318.38 ms; SD = 13.99 ms). Task performance was modulated by the experimental manipulation of stimulus certainty. As expected, accuracy was highest and reaction times were fastest for stimuli far from the category boundary (High-Certainty) and decreased for stimuli close to the boundary (Low-Certainty) (Figure 3). All the following analyses were performed using correct trials only.

**Figure 3.**
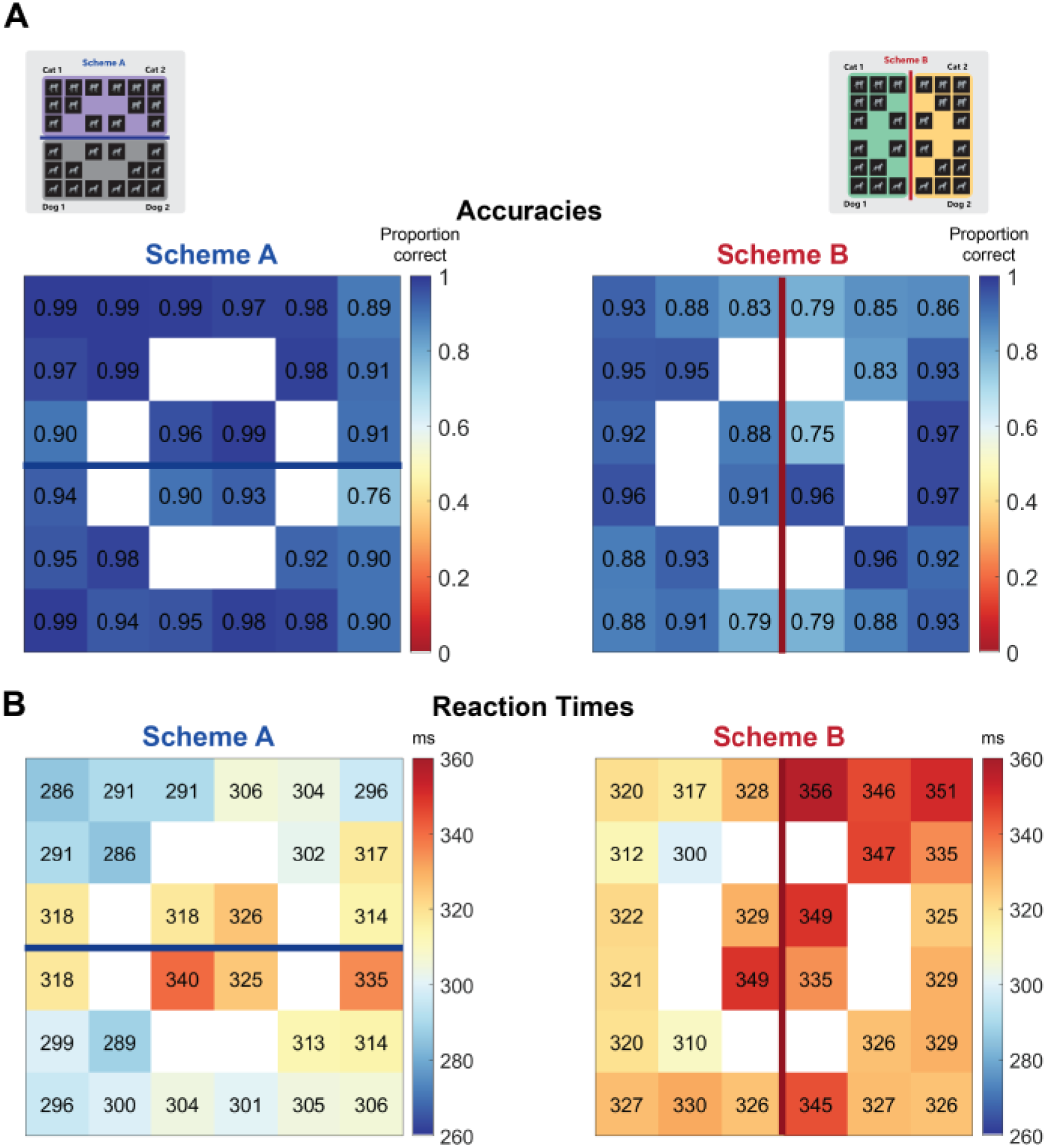
Average **(A)** Accuracy and **(B)** Reaction Time of monkeys in each stimulus and scheme condition. Accuracy is shown as proportion of trials with a correct response and Reaction time as the time between the test stimulus onset and response.

### Coding of Scheme and Stimulus Information in IT and PFC

Our first analysis of the neural data looked at the neural representations in IT and PFC at population level to characterize the temporal dynamics of task and stimulus information coding using different lenses as defined by our hypothesized models (Figure 4).

**Figure 4.**
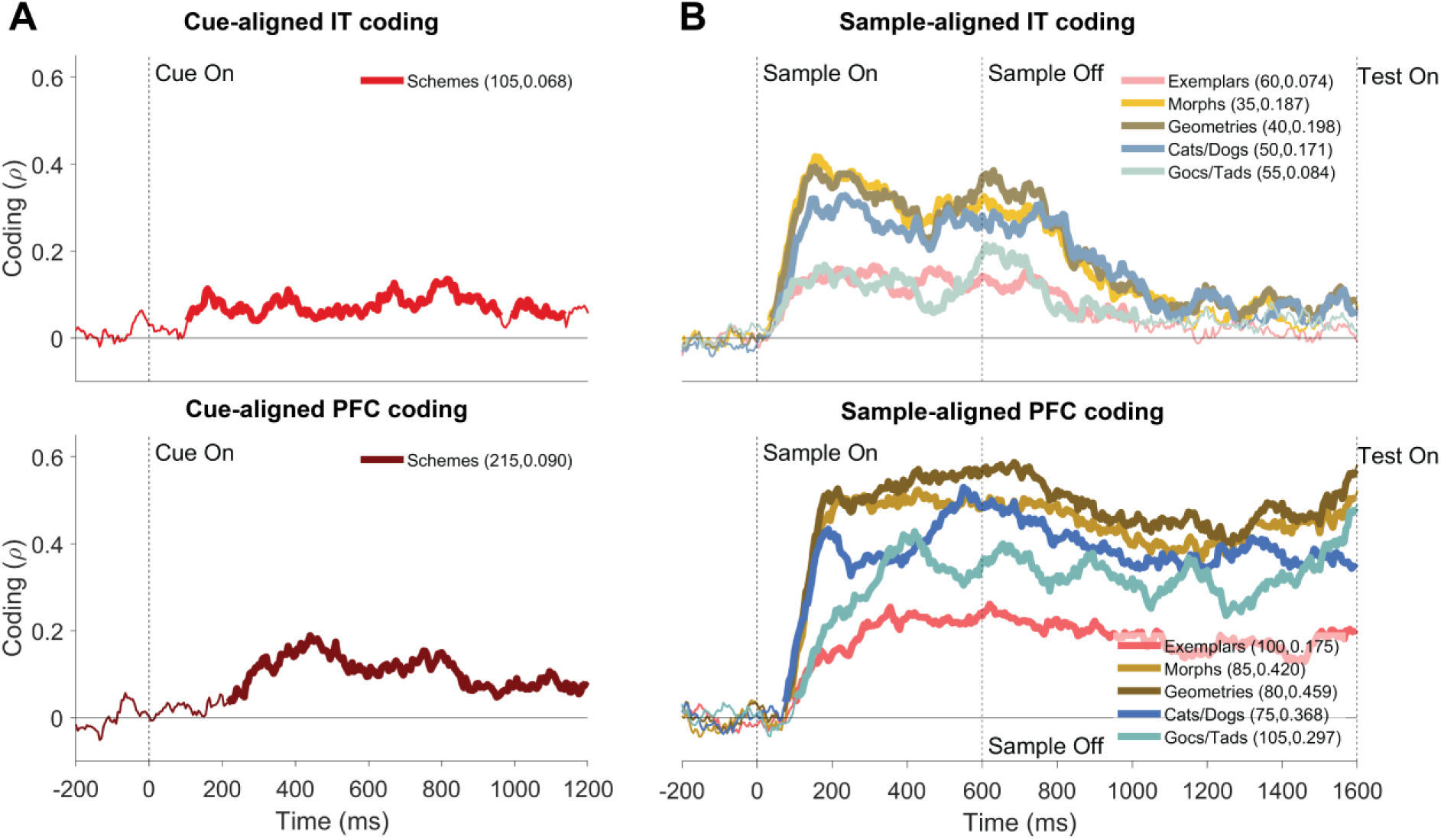
Population-level scheme and stimulus coding. **(A)** Cue-aligned scheme information coding in IT (top) and PFC (bottom). **(B)** Sample-aligned stimulus information coding in IT (top) and PFC (bottom). Different line colors correspond to different conceptual models. The values in parentheses show the onset time of significant neural coding in milliseconds (left number) and the average post-onset area-under-the-curve (AUC, right number). Thickened lines indicate the time points when information coding was significantly above chance (permutation test, p < 0.05; cluster-corrected for FWER). Vertical dashed lines mark critical trial events (i.e., cue, sample onset and offset).

Following the cue onset, scheme information emerged earlier in IT (105 ms) than in PFC (215 ms). This was predictable and can be explained by the precedence of coding of visual differences in the cues in IT possibly followed by the extraction of both visual and scheme information in PFC. Monkeys had to rely on the visual cues to extract the scheme information and the first bump of information in PFC (∼450ms) likely reflects the scheme information extracted from visual cues. Both areas maintained the scheme information until the sample stimulus came on the screen (∼1s later), which was when the monkeys needed to do the categorization.

To see how abstract the stimulus representations in IT and PFC were, we examined the neural representations during the sample presentation and delay using five stimulus-based models. These models detected different levels of abstraction from concrete stimulus-specific encoding to abstract categorical distinctions (defined in Figure 2A; for the results of all models see Figure S1). The order of models which quantified different levels of abstraction were similar across IT and PFC, with more robust information for semi-abstract compared to concrete or abstract representations (Figure 4). Specifically, in IT, information coding was earlier and stronger for the Morphs (35ms, AUC=0.187) and Geometries (40ms, AUC=0.198) models than Cats/Dogs (50ms, AUC=0.171), Gocs/Tads (55ms, AUC=0.084) and Exemplars (60ms, AUC=0.074) models. Similarly in PFC, information coding was earlier and stronger for Morphs (85ms, AUC=0.420) and Geometries (80ms, AUC=0.459) models than Cats/Dogs (75ms, AUC=0.368), Gocs/Tads (105ms, AUC=0.297) and Exemplars (100ms, AUC=0.175) models. These suggested that rather than a pure low-level image-level representation of exemplars or an abstract category-level summary representation, the two areas maintained a relative map of the relationship between stimuli, which was semi-abstract, as captured by Morphs and Geometries models.

The observed difference in informational magnitude (AUC) between the cue-aligned (schemes information) and sample-aligned (stimulus information) periods reflects a transition from the sparse encoding of abstract, low-energy task-rule identifiers (the category scheme) to the robust, high-magnitude population responses elicited by high-contrast, complex visual stimuli during the sample period.

IT consistently exhibited earlier information coding across all tested representational models compared to PFC. While IT coding was rapid and transient, decaying shortly after stimulus offset, PFC representations were more sustained, persisting throughout the entire memory delay period (Figure 4B). This established a hierarchy where IT provides early, transient sensory evidence, and PFC maintains the task-relevant information. These results were not explained by higher number of neurons recorded in PFC than IT as a similar pattern was observed when subsampling the PFC neurons to the same number of neurons in IT (Figure S2).

### More Sustained Recurrence of Information in PFC than IT

The above coding profiles suggested more sustained coding of scheme and stimulus-related information in PFC than IT. To characterize how sustained the information coding was over time, we developed a model-based method to evaluate how past representations in an area predicted present representations in the same area reflecting information recurrence (Eq. 1). This method, which extends previous time-generalization methods (King & Dehaene, 2014) to non-binary discriminations, allowed us to evaluate the recurrence of distinct types of information using our hypothesized models.

Consistent with expectations from hierarchical processing and recurrent integration theories (Buschman & Kastner, 2015; Freedman et al., 2003), the temporal profile of recurrence differed markedly between IT and PFC (Figure 5). For most types of information, recurrence in IT emerged earlier and showed a relatively shorter duration than PFC typically fading out during the memory delay period. In contrast, PFC recurrence tended to start later but persisted throughout the stimulus and delay periods, indicating sustained internal reactivation likely linked to working memory or information maintenance (Fuster, 2001; Miller et al., 2018).

**Figure 5.**
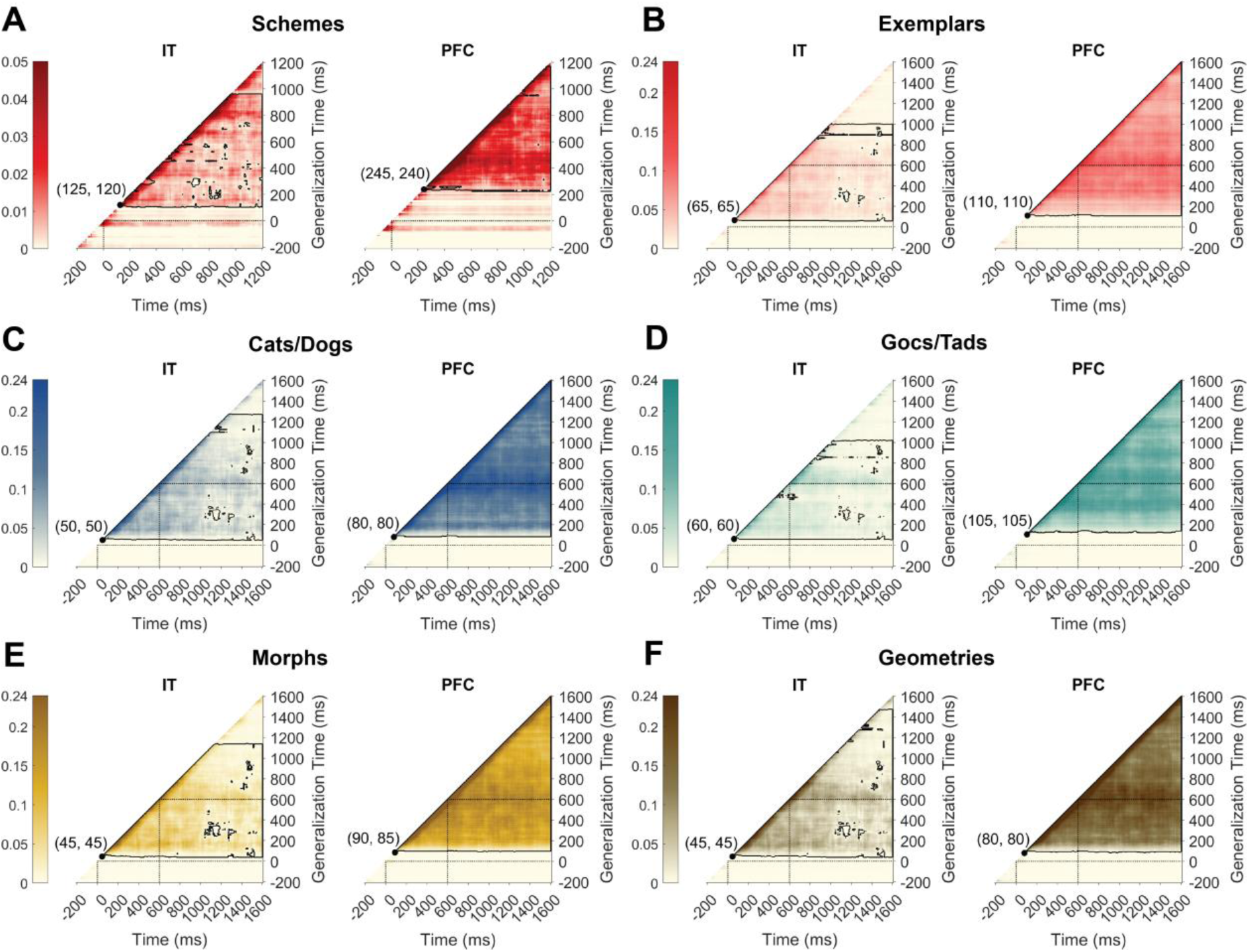
Recurrence of distinct types of information in IT and PFC. Each color indicates a different type of information as defined by a different model: **(A)** Schemes, **(B)** Exemplars, **(C)** Cats/Dogs, **(D)** Gocs/Tads, **(E)** Morphs, and **(F)** Geometries. X and y-axes show the present and generalization time, respectively. Generalization refers to the past time when the information pattern is borrowed from to predict the present-time information. Black contours enclose significant recurrence (permutation test, p < 0.05; cluster-corrected for FWER). Left and right numbers in parentheses indicate the first significant recurrence on the time and generalization axes, respectively. Dashed lines mark critical events in the trial.

Among stimulus- and task-based models, IT exhibited earliest recurrence for the Morphs and Geometries models, both appearing at approximately 45 ms post-stimulus onset. Notably, Geometries recurrence in IT was not only early but also the most prolonged, suggesting stable internal reprocessing of relational exemplars information. In contrast, the Scheme model, which captured information about the categorization scheme/task, showed the latest recurrence onset in IT (125 ms), while Exemplars, probably the more concrete stimulus-based model, followed with a moderate delay (65 ms). In the PFC, recurrence also emerged earliest for the Geometries and Cats/Dogs models (85 ms), consistent with early integration of relational and categorical structure. Recurrence for the Schemes model was significantly delayed (245 ms), and Exemplars emerged at 110 ms, reinforcing the idea that abstract and behaviorally salient features dominate early recurrence dynamics in PFC, whereas concrete representations take longer to reappear or may be deprioritized in ongoing cognitive processing.

### Exchange of Task and Stimulus Information between IT and PFC

To provide a new lens onto the IT-PFC interactions during categorization, we analyzed the dynamic flow of information between IT and PFC using our novel approach of RCA (Karimi-Rouzbahani et al., 2021, 2022). RCA allowed us to determine the flow of distinct types of information between these areas using the different conceptual models defined above. Consistent with the hierarchical organization of visual-cognitive pathways, feed-forward (FF) flow was defined as the directional influence of past IT representations on present PFC representations, reflecting bottom-up propagation of dominantly sensory information. Conversely, feedback (FB) flow was defined as the top-down modulation of present IT representations by past PFC representations, reflecting higher-order task-dependent contextualization of sensory information. We also provide information coding as the top panel to allow for cross-checking of information flow and coding.

During the cue period, there was a significant FF flow of task (Schemes) information from IT to PFC (Figure 6A) shortly after the cue onset (105ms). This initial bottom-up transmission was followed by feedback flow starting at 220ms with longer delays. The FF and FB flows showed a cyclical exchange between IT and PFC in the following windows. This bidirectional circulation suggests recurrent processing, where early sensory evidence (e.g., cue information) is progressively integrated with top-down task-related (scheme) information.

**Figure 6.**
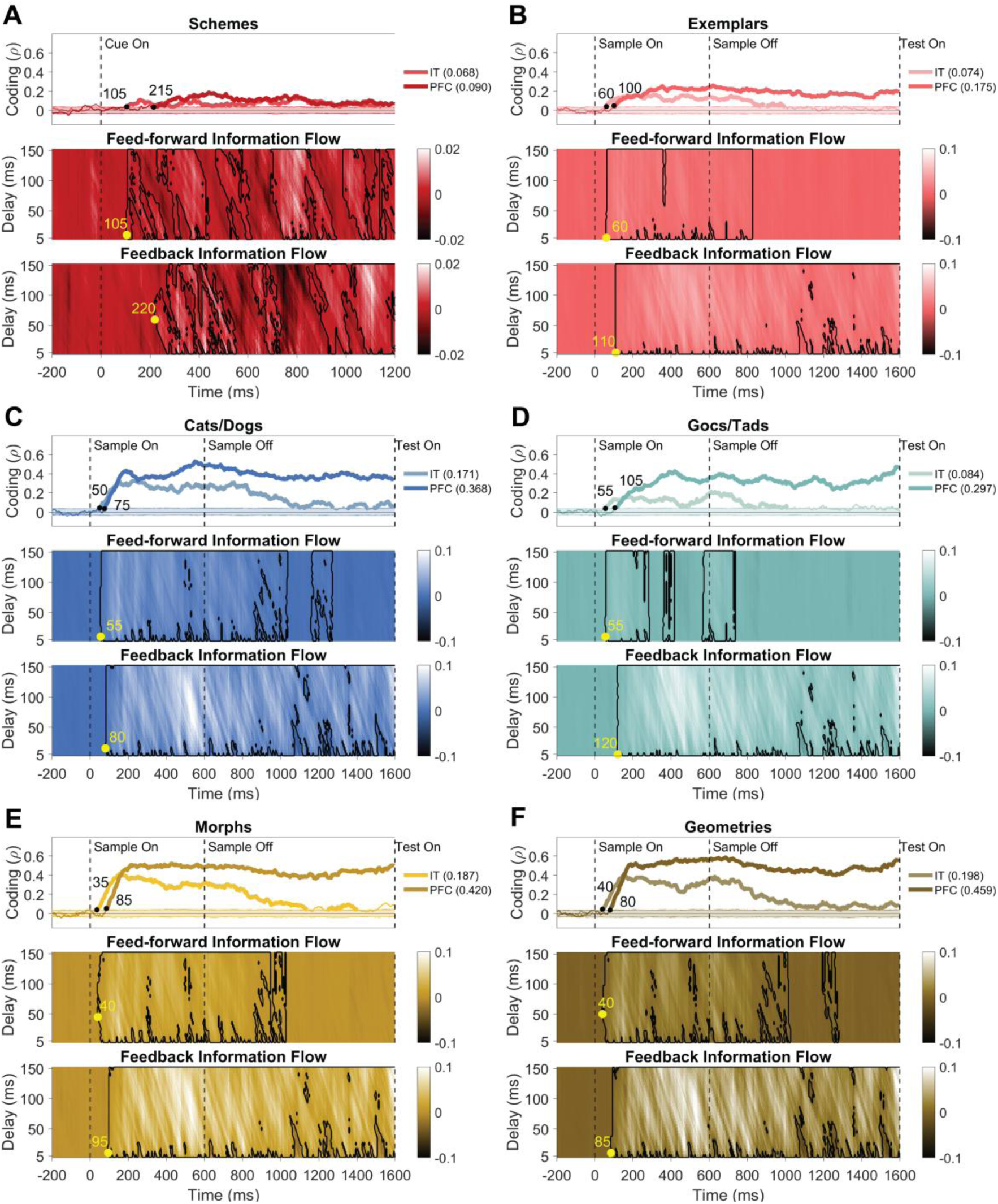
Flows of different types of information between IT and PFC. Each color indicates a different type of information as defined by a different model: **(A)** Schemes, **(B)** Exemplars, **(C)** Cats/Dogs, **(D)** Gocs/Tads, **(E)** Morphs, and **(F)** Geometries. Each panel contains three plots. **Top** (Information Coding) in IT (light color) and PFC (dark color). Thickened lines indicate time points with significantly above-chance coding (permutation test, p < 0.05; cluster-corrected for FWER) and the around-zero shadings reflect the null coding distribution obtained by shuffling the condition labels 1000 times. The dots and corresponding numbers show the time point of the first significant coding. **Middle** (Feed-Forward flow): IT-to-PFC information flow over time (x-axis) and transmission delays (y-axis: 5-150 ms). Black contours enclose significant flows (permutation test, p < 0.05; cluster-corrected for FWER), with yellow dots marking the first significant latencies (values in ms). **Bottom** (Feedback flow): PFC-to-IT information flow, analyzed identically to feed-forward flow, but in the opposite direction.

During the sample period, information flows for both abstract and concrete stimulus-based models followed a relatively similar pattern, with earlier FF followed by FB flows (Figure 6B-F). The FF and FB flows were both active during the stimulus presentation period supporting recurrent processing of information between IT and prefrontal cortices during visual processing (Kar & DiCarlo, 2021; Noroozi et al., 2024). However, while FF flow from IT to PFC terminated shortly after the stimulus offset, FB flows persisted during the memory delay period. The continued FB flow after the stimulus disappearance (despite FF termination) implies that PFC transitions from a receiver to a transmitter of information, potentially facilitating stimulus information retrieval in sensory areas upon the presence of test image (Dijkstra et al., 2017; S.-H. Lee & Baker, 2016).

### Dynamic Reconfiguration of Information Flow by Stimulus Certainty

The most critical question of this study was whether stimulus uncertainty modulated information flow between sensory and higher order cognitive areas. To address this, we separately quantified the coding and flow of information for stimuli far from and close to the categorical boundaries using “Far from Boundary” and “Close to Boundary” models, respectively (Figure 2A; called Far and Close here for brevity). This distinction can be important as this can test two alternative hypotheses: stimuli closer to the boundary are prone to lead to behavioral errors in the categorization task. Therefore, they might be represented more strongly than stimuli far from the boundary. Alternatively, as distant stimuli are visually more distinct, they might be represented more strongly than close-to-boundary stimuli which are visually less discriminable. Importantly, we have shown in the past that that the uncertainty in the sensory stimuli (i.e., the signal to noise ratio) can modulate the level of FF and FB flows in the brain (Karimi-Rouzbahani et al., 2021) and would like to test if this remains the case when perceptual uncertainty was imposed by stimulus similarities across categories rather than added noise.

Far stimulus information coding started in the IT at approximately 40 ms post-stimulus onset followed by PFC engagement at 85 ms (Figure 7A upper panel). While IT representations were more transient, PFC representations showed the canonical sustained representations maintained after the stimulus offset.

**Figure 7.**
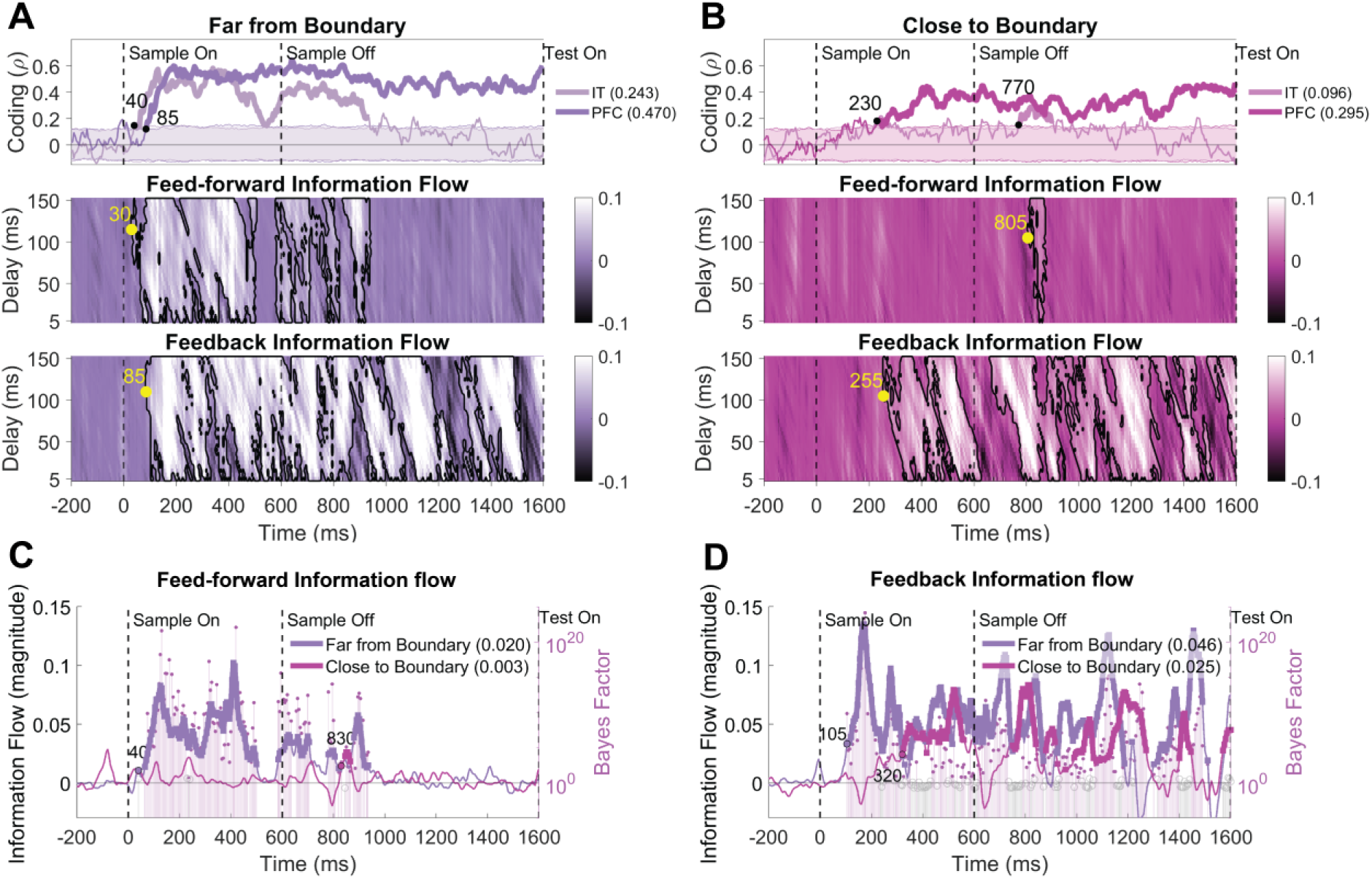
Modulation of coding and flows of stimulus information between IT and PFC with stimulus certainty. **(A)** Far from Boundary and **(B)** Close to Boundary stimulus information. **Top** (Information Coding) in IT (light color) and PFC (dark color). Thickened lines indicate time points with significantly above-chance coding (permutation test, p < 0.05; cluster-corrected for FWER) and the around-zero shadings reflect the null coding distribution obtained by shuffling the condition labels 1000 times. The dots and corresponding numbers show the time point of the first significant coding. **Middle** (Feed-Forward flow): IT-to-PFC information flow over time (x-axis) and transmission delays (y-axis: 5-150 ms). Black contours enclose significant flows (permutation test, p < 0.05; cluster-corrected for FWER), with yellow dots marking the first significant latencies (values in ms). **Bottom** (Feedback flow): PFC-to-IT information flow, analyzed identically to feed-forward flow, but in the opposite direction. **(C)** Delay-averaged feed-forward and **(D)** feedback flows for Far (purple) and Close (pink) boundary conditions. Thickened lines indicate timepoints which were significant across >50% of delays. Vertical bars are Bayes Factor (BF) values where filled circles show evidence for difference (BF > 6) and empty gray circles show insufficient evidence (BF < 6) for difference between the two flows. Significantly above-chance flow onset is indicated by the black filled circles with ms labels. The average post-onset AUC for each model is indicated in the parentheses. Vertical dashed lines refer to sample onset and offset.

Far stimulus information flow mirrored patterns observed in earlier stimulus models: initial FF flow from IT emerged during early stimulus presentation (30 ms), followed by the onset of FB flow from PFC beginning around 85 ms (Figure 7A, middle panel: FF and lower panel: FB). Notably, while FF transmission was transiently suspended after stimulus offset, FB flow from PFC to IT persisted, consistent with trends observed across other stimulus models.

In contrast to the Far stimulus information, the Close stimulus information exhibited an inversion of the expected cortical processing sequence. Neural coding did not initiate in the IT; instead, representations first emerged in the PFC after a prolonged delay of approximately 230 ms (Figure 7B, upper panel). Throughout the stimulus presentation, no significant Close stimulus information was detected in IT (p > 0.05, cluster-corrected), indicating a lack of bottom-up encoding of close-to-boundary stimuli under high uncertainty. Interestingly, IT engagement only became evident during the memory delay period, with a latency of 550 ms relative to the initial PFC representations.

Information flow analyses revealed a pattern consistent with this reordering in information coding dynamics. FB flow from PFC to IT began around 255 ms post-stimulus onset and persisted intermittently throughout the stimulus presentation, ceasing near stimulus offset (Fig 7B, middle panel). Notably, approximately 100 ms after stimulus offset, FB flow sharply increased, potentially reflecting a post-hoc reactivation of IT for reevaluation or delayed encoding (Fig 7B, lower panel). This surge was shortly followed by a brief appearance of FF flow from IT to PFC at around 805 ms, suggesting a delayed and possibly reconstructed transfer of stimulus information toward decision-related regions.

To directly and statistically compare the magnitudes of FF and FB flows for the Far and Close information, we averaged information flows across all delays (5-150 ms), yielding delay-averaged FF and FB flows shown in Figure 7C and 7D. As demonstrated in Figure 7C, FF flow was stronger (BF > 6) for Far stimuli compared to Close stimuli in majority of time points during the stimulus presentation and delay period (Far vs. Close AUCs= 0.020 vs. 0.003). There was also stronger (BF > 6) and more FB flows of information (Figure 7D) for the Far than Close stimulus information both during and after stimulus presentation (Far vs. Close AUCs= 0.046 vs. 0.025). These results suggest that while Far stimulus information became available in IT and propagated to PFC before being circulated between them, Close stimulus information first appeared in PFC and was fed back to IT before circulation. This supports the role of PFC in enhancing sensory representations in lower sensory-involved areas (Kar & DiCarlo, 2021; Noroozi et al., 2024).

### Improved Information Coding and Flow for Task-Relevant Categories

We then asked whether behavioral relevance of stimulus category enhanced stimulus representation. This is important for a cognitive task that relevant aspects of the task are prioritized over irrelevant aspects so that the cognitive resources are focused on the relevant information for task performance. To test this, Relevant and Irrelevant Categories models were fitted to the neural data and compared. As previous studies have shown, stimulus distinctions which matched the cued categorization scheme were encoded more strongly in the human brain as they match the behavioral needs (Duan et al., 2024; J. Jackson et al., 2017; J. B. Jackson et al., 2021). We were wondering how this would be represented in the monkey brain and if task relevance enhanced the flow of information between IT and PFC.

For both Relevant and Irrelevant Categories, the information coding appeared in IT before PFC, with the characteristic temporal pattern of transient coding in IT and sustained coding in PFC throughout stimulus presentation and delay periods (Figure 8A, top panels). However, critical differences emerged in both timing and magnitude. Relevant category coding in IT reached significance earlier compared to Irrelevant Categories (40 ms vs. 75 ms). Moreover, Relevant Categories showed a higher coding throughout the analysis windows than Irrelevant Categories (IT: 0.094 vs. 0.074 and PFC: 0.260 vs. 0.204). These supported an advantage in the coding of scheme-matching categories than those orthogonal to the schemes and behavior.

**Figure 8.**
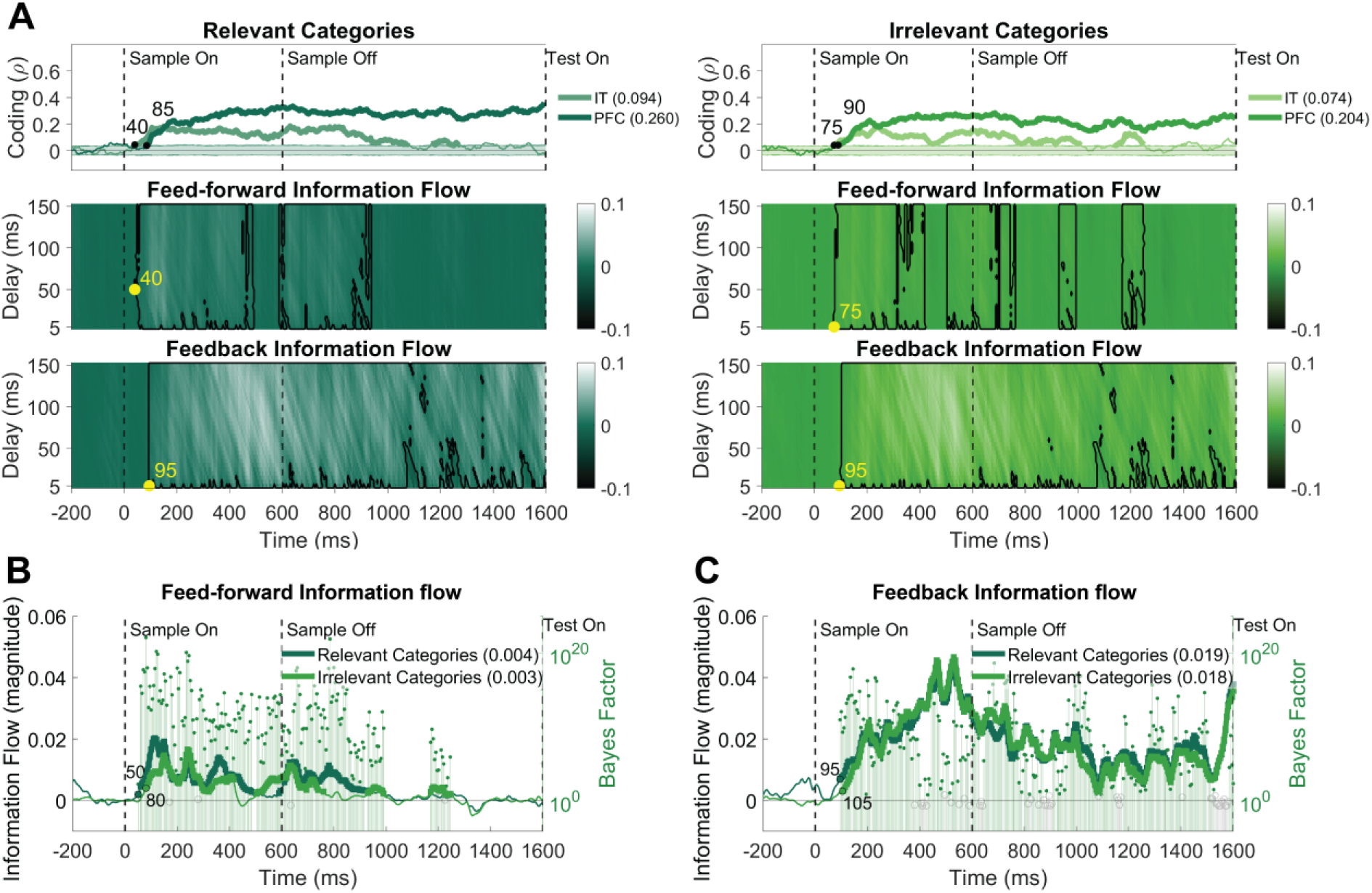
Modulation of coding and flows of stimulus information between IT and PFC with scheme relevance. **(A)** Left: Relevant, Right: Irrelevant stimulus information. **Top** (Information Coding) in IT (light color) and PFC (dark color). Thickened lines indicate time points with significantly above-chance coding (permutation test, p < 0.05; cluster-corrected for FWER) and the around-zero shadings reflect the null coding distribution obtained by shuffling the condition labels 1000 times. The dots and corresponding numbers show the time point of the first significant coding. **Middle** (Feed-Forward flow): IT-to-PFC information flow over time (x-axis) and transmission delays (y-axis: 5-150 ms). Black contours enclose significant flows (permutation test, p < 0.05; cluster-corrected for FWER), with yellow dots marking the first significant latencies (values in ms). **Bottom** (Feedback flow): PFC-to-IT information flow, analyzed identically to feed-forward flow, but in the opposite direction. **(B)** Delay-averaged feed-forward and **(C)** feedback flows for Relevant (dark green) and Irrelevant (light green) stimulus conditions. Thickened lines indicate timepoints which were significant across >50% of delays. Vertical bars are Bayes Factor (BF) values where filled circles show evidence for difference (BF > 6) and empty gray circles insufficient evidence (BF < 6) for difference between the two flows. Significantly above-chance flows are indicated by the black filled circles with ms labels. The average post-onset AUC for each model is indicated in the parentheses. Vertical dashed lines refer to sample onset and offset.

Next, we looked at the information flows. Relevant and Irrelevant Category information flowed in FF direction before FB flows started as was the case for other types of information (Figure 8A, middle panels). Interestingly, following the advantage in their coding, Relevant Category information started flowing in the FF direction earlier (40 ms vs. 75 ms), and showed longer periods of sustained flow than the Irrelevant Category flow, which showed a patchy pattern of FF flows. Evaluation of FB flows for the Relevant and Irrelevant Category information showed relatively similar patterns (Figure 8A, bottom panels). Direct and statistical comparison of delay-averaged FF and FB flows of Relevant and Irrelevant Categories also showed similar results (Figure 8B-C). We observed systematically earlier (50 ms vs. 80 ms) and stronger (AUCs = 0.004 vs. 0.003) FF flow for Relevant vs. Irrelevant Category information (Figure 8B). This was especially evident in the earlier windows of time after the stimulus onset before the 150 ms. The advantage of category relevance was less obvious in FB flows with some advantage in temporal precedence (95 ms vs. 105 ms) and magnitudes (AUCs= 0.019 vs. 0.018) of FB flows for the Relevant vs. Irrelevant Category information (Figure 8C).

### Stimulus Certainty Modulates Behavioral Information Coding and Flows

Next, we tested to see if behavioral performance was reflected in neural representations. To that end, we constructed a Reaction Time representational dissimilarity matrix (RDM) that quantified pairwise differences in RTs across experimental conditions. This RT RDM served as a model to probe the neural representation of reaction time: if a brain region exhibits representations that correlate with the RT model, that region is involved in behavioral performance. The RT model looks for behavioral information beyond perceptual or conceptual similarity between conditions.

Analyzing the response-aligned data showed that reaction times were represented in PFC earlier than in IT, particularly in the final moments leading up to the motor response (Figure 9). Notably, significant RT-related coding in PFC emerged approximately 400 ms before the behavioral response—a timepoint that proceeds the maximum observed reaction time from the two monkeys, which was 359 ms. This early coding of RT might reflect a proactive adjustment in cognitive control, such as response readiness, attentional allocation, or decision threshold setting which varies in a trial-wise manner and controls response generation. Such preparatory dynamics are consistent with prior work showing internally driven fluctuations in PFC prior to stimulus onset that predict behavioral variability (Miller & Cohen, 2001). IT region reflected the behavioral RTs closer to the response (starting at -270ms), possibly reflecting late-stage sensory evaluation or outcome-related feedback processing. These findings might highlight a top-down anticipatory component of behavioral performance, suggesting that PFC dynamically configures task-state parameters that influence downstream processing and response latency.

**Figure 9.**
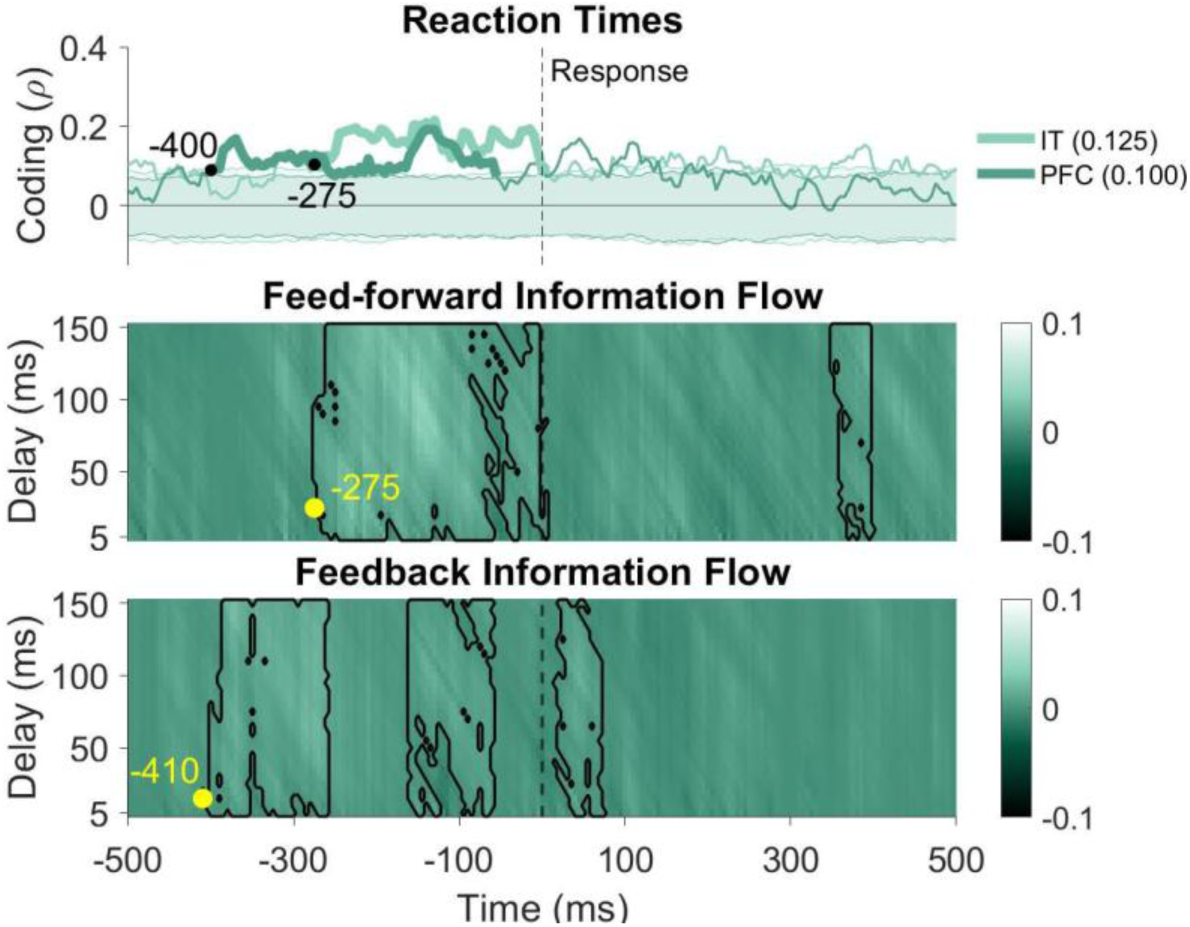
Neural correlates of behavioral performance. Response aligned coding and flow of reaction time information. **Top** (Information Coding) in IT (light color) and PFC (dark color). Thickened lines indicate time points with significantly above-chance coding (permutation test, p < 0.05; cluster-corrected for FWER) and the around-zero shadings reflect the null coding distribution obtained by shuffling the condition labels 1000 times. The average pre-response AUC for each model is indicated in the parentheses. **Middle** (Feed-Forward flow): IT-to-PFC information flow over time (x-axis) and transmission delays (y-axis: 5-150 ms). Black contours enclose significant flows (permutation test, p < 0.05; cluster-corrected for FWER), with yellow dots marking the first significant latencies (values in ms). **Bottom** (Feedback flow): PFC-to-IT information flow, analyzed identically to feed-forward flow, but in the opposite direction. The dots and corresponding numbers show the time point of the first significant coding or flow. Vertical dashed lines refer to response onset.

Information flow analyses revealed a pattern consistent with the order of information coding in IT and PFC (Figure 9). FB flow from PFC to IT began around -410 ms before the response while the FF flow from IT to PFC followed at -275 ms before the response. This suggests the initial processing of behavioral information in PFC which is propagated to IT. This supports the dominant processing of behavioral information in higher-order frontal brain areas.

Finally, to determine if task demand influences inter-areal communication, we analyzed information flow using models combining Reaction Time (RT) and Perceptual Certainty (Figure 10). Behavioral information coding was robustly present in both areas, with higher magnitudes for “Close” versus “Far” stimuli (IT AUC: 0.129 vs. 0.177; PFC AUC: 0.289 vs. 0.403). Crucially, response-aligned information transfer was initiated earlier for uncertain (“Close”) trials. PFC initiated RT-related feedback at -370ms in the Close condition, compared to -315ms for Far stimuli. Similarly, IT onsets shifted from -250ms (Close) to -110ms (Far). Although feedback (PFC→IT) preceded feedforward flow in both cases, the accelerated onset under uncertainty suggests that the top-down recruitment of behavioral parameters was prioritized when sensory evidence was ambiguous.

**Figure 10.**
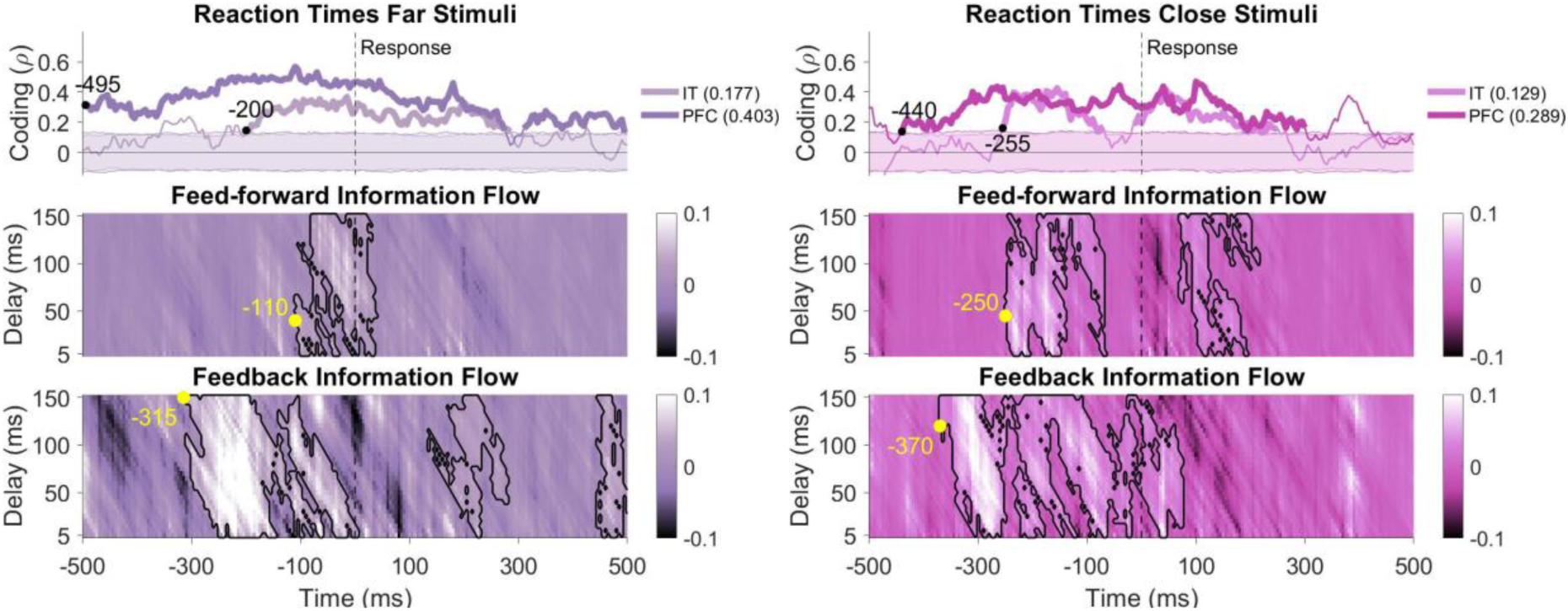
Stimulus Certainty Modulates Behavioral Information Coding and Flow. Response aligned coding and flow of reaction time information as a function of perceptual certainty (left: RT model × Far model; right: RT model × Close model). **Top** (Information Coding) in IT (light color) and PFC (dark color). Thickened lines indicate time points with significantly above-chance coding (permutation test, p < 0.05; cluster-corrected for FWER) and the around-zero shadings reflect the null coding distribution obtained by shuffling the condition labels 1000 times. The average pre-response AUC for each model is indicated in the parentheses. **Middle** (Feed-Forward flow): IT-to-PFC information flow over time (x-axis) and transmission delays (y-axis: 5-150 ms). Black contours enclose significant flows (permutation test, p < 0.05; cluster-corrected for FWER), with yellow dots marking the first significant latencies (values in ms). **Bottom** (Feedback flow): PFC-to-IT information flow, analyzed identically to feed-forward flow, but in the opposite direction. The dots and corresponding numbers show the time point of the first significant coding or flow. Vertical dashed lines refer to response onset.

## Discussion

Our study provides a tripartite information-based lens, through the dynamics of *information coding*, *recurrence*, and *inter-areal flow*, to dissect the distinct yet interactive roles of the inferotemporal (IT) and prefrontal cortex (PFC) in visual categorization. We examined how task- and stimulus-related information was encoded, dynamically modulated, and transferred across IT and PFC along different category schemes. By applying representational models of varying abstraction, task relevance, and stimulus certainty, we provided a detailed account of when, where, and how categorical information emerged in neural population activity.

Our **information coding** results showed that IT consistently exhibited earlier coding than PFC, aligning with its role as a fast visual processing hub (DiCarlo et al., 2012; Muhammad et al., 2006). Moreover, our analysis revealed that relational semi-abstract models—capturing stimulus similarities through morphometric and geometric structure—elicited stronger population-level coding than either concrete exemplar-level or abstract category-level distinctions. While these models align with theoretical works describing boundary-based models(Ashby & Gott, 1988; Ashby & Maddox, 1992), we refer to them as relational exemplar representations. This distinction is critical: unlike traditional models that emphasize a fixed decision surface, our models capture the topological continuity among exemplars, occupying a unique middle ground between concrete encoding and abstract rule-based classification (Barrett & Miller, 2026; Minda et al., 2024). Quantitatively, this relational coding was the strongest neural representation in both areas, evidenced by significantly higher coding strength (e.g., AUCs of Morphs: 0.187 in IT, 0.420 in PFC) compared to stimulus-specific (Exemplars: 0.074 in IT, 0.175 in PFC) or abstract models (Categories: 0.171 in IT, 0.368 in PFC). This robust encoding suggests that both IT and PFC are optimized for building a continuous, topological map of object feature space to facilitate generalization (Baldassi et al., 2013; Tanaka, 1996), rather than relying on highly specific ’grandmother-cell’ representations (Gross, 2002). The delayed emergence of abstract category information in IT further suggests these formats are computed from this underlying relational structure.

The temporal dynamics, evaluated through **recurrence analysis**, revealed a clear functional distinction. While IT consistently exhibited the earliest coding onset across almost all tested models, serving as the primary source of bottom-up information, this information was highly transient, decaying rapidly after stimulus offset. The recurrence analysis confirmed this transient nature, showing that IT’s internal reprocessing of information was short-lived (Brincat & Connor, 2004; DiCarlo et al., 2012). In contrast, the PFC demonstrated later information onset but sustained coding and active recurrence throughout the delay period. This sustained activity is a hallmark of PFC function, widely linked to working memory and rule maintenance (Gilbert & Li, 2013; Miller & Cohen, 2001). Our recurrence analysis builds on previous studies by showing that PFC does not just passively remain active (Funahashi et al., 1989; Fuster & Alexander, 1971); it actively and persistently reactivates and reprocesses task information internally. This content-specific recurrence confirms the PFC’s role in maintaining the high-level category rule required for the task (Gilbert & Li, 2013).

Complementing these coding and recurrence analyses, our initial **information flow** analyses revealed a robust temporal cascade of feed-forward (FF) followed by feedback (FB) information flow between IT and PFC. Across both task-based and stimulus-based models, initial FF flow from IT to PFC during early stimulus presentation was followed by delayed FB flow from PFC back to IT. This cyclical exchange is consistent with recurrent models of perception and decision-making, where bottom-up signals are iteratively integrated with top-down contextual or rule-based information (Bastos et al., 2012; Friston, 2005). It is also consistent with monkey studies which used causal suppression techniques, to show that inactivation of ventrolateral PFC can impact visual representations in IT (Kar et al., 2019; Kar & DiCarlo, 2021; Debes & Dragoi, 2023). After stimulus offset, FF transmission ceased while FB flow persisted, indicating that PFC sustains and projects information back to IT, potentially to support working memory maintenance or predictive processing during memory delays (Miller et al., 2018; Tomita et al., 1999). This established IT - > PFC -> IT cycle serves as the typical flow profile under high-certainty conditions.

Our most significant finding is that the direction of information flow between IT and PFC is not fixed but is dynamically reconfigured by the perceptual certainty of the stimulus. This modulation was observed precisely where top-down control is predicted to be most critical: in conditions of sensory ambiguity. For high-certainty (Far-from-boundary) stimuli, the canonical FF flow dominated early in the trial. Crucially, uncertain (Close-to-boundary) stimuli triggered a dominant, reordering of the cortical processing sequence: PFC encoded category information first, and a rapid feedback flow (PFC -> IT) emerged, preceding the canonical feed-forward sweep. This dynamic reconfiguration of IT-PFC hierarchy supports theories of flexible cortical hierarchy and aligns with predictions from the Reverse Hierarchy Theory (RHT) (Ahissar & Hochstein, 2004), which posits that difficult perception is primarily achieved via top-down processing from executive areas. Our results extend this idea by showing that the ‘reconfiguration’ is not just slower access, but an active, content-specific change in the dominant information transfer direction, mediated by the categorical information necessary to resolve the ambiguity. This reconfiguration mechanism is also consistent with the computational goals of the Hierarchical Predictive Coding (HPC) framework. HPC posits that the brain continuously generates top-down predictions to minimize prediction error (PE) signaled by bottom-up inputs (Friston, 2005). In our context, sensory ambiguity (Close-to-boundary stimuli) would generate a substantial and persistent PE in the ventral visual stream (IT) because the sensory features conflict with initial high-level category expectations. The observed PFC->IT flow thus provides a precise, neurophysiological mechanism for PE minimization: the PFC transmits a high-level prior or prediction about the most likely category to IT. This top-down signal, or a signal increasing its computational weight (Precision), effectively suppresses the noisy, unreliable bottom-up PE signal in IT, thereby resolving the sensory conflict and allowing the categorical decision to stabilize. This interpretation links the temporal dynamics of RHT with the computational necessity of Bayesian inference, offering a unified explanation for the role of executive feedback in uncertain perception. Our behavioral-neural correlate analysis further reinforces this functional link; we observed that information regarding the upcoming reaction time (RT) is transmitted from the PFC to IT before being consolidated in the final prefrontal decision state. This suggests that the PFC-to-IT feedback is not merely a passive echo, but a ’behavioral priming’ signal that shapes sensory processing to meet the temporal demands of the task. This finding provides a concrete behavioral anchor for our ’resolution mode’ hypothesis, showing that the top-down reordering of the hierarchy is directly coupled to the animal’s decision-making process.

When interpreting the earlier emergence of feedback under uncertainty, we considered whether the smaller IT population (n=154) compared to the PFC (n=549) might bias our results toward a ’late’ IT onset. While our IT sample size is consistent with established primate electrophysiology studies in this domain (Kar & DiCarlo, 2021; Majaj et al., 2015; Noroozi et al., 2024), we acknowledge that a larger population might reveal lower-magnitude signals currently below our detection threshold. However, our internal controls suggest that the observed temporal sequence is not a byproduct of sample size. Subsampling controls further demonstrate that these temporal dynamics are independent of neuron count (Figure S2).

Specifically, when equalizing population sizes (n=154), the qualitative hierarchy reversal persisted: feedforward flow dominated the ’Far’ condition, while it was entirely suppressed during the stimulus window for ’Close’ stimuli, with feedback emerging as the primary signal during the delay (Figure S3). This confirms that the absence of early feedforward drive under uncertainty is a genuine physiological consequence of stimulus ambiguity rather than an artifact of sampling power. Beyond population size, we also considered whether a global reduction in signal-to-noise ratio (SNR) due to task difficulty could explain these shifts. Critically, a purely SNR-based model predicts a quantitative delay in signal emergence while maintaining the canonical IT-to-PFC feedforward sequence. In contrast, our data reveals qualitative temporal reordering, where PFC leads IT by several hundred milliseconds for ambiguous stimuli. This is further corroborated by our analysis of combined behavioral-perceptual models (Figure 10), which shows that behavioral information flow is initiated significantly earlier for uncertain versus certain stimuli. The observation that the brain initiates this content-specific transfer earlier when stimuli are ambiguous—despite a ‘weaker’ sensory signal—strongly argues for a proactive recruitment of top-down assistance rather than a passive SNR-driven delay. This shift aligns with the ‘Challenge Stimulus’ framework (Kar et al., 2019; Kar & DiCarlo, 2021) and the consensus that ‘hard’ visual tasks require late-stage recurrent signals and top-down intervention (Fyall et al., 2017; Kang et al., 2026; Kreiman & Serre, 2020; Namima & Pasupathy, 2021; Noroozi et al., 2024; Rajaei et al., 2019; Tang et al., 2018). By showing that PFC leads IT under uncertainty, our RCA framework provides a specific directional mechanism for the recurrent computations often observed in these complex categorization tasks.

The context-dependent reordering of information flow observed here challenges the classical view of a fixed, feedforward (FF) hierarchy (Felleman & Van Essen, 1991), suggesting instead that the brain leverages its bidirectional anatomy as a ’heterarchy’ (Pessoa, 2023). In this framework, the PFC serves as a dynamic regulatory hub that selectively gates top-down influence, enforcing a dominant PFC-to-IT flow only when task demands require executive intervention to resolve perceptual uncertainty. Importantly, while we interpret this reconfiguration through the lens of cortico-cortical recurrence, the potential role of subcortical and transthalamic pathways, such as the mediodorsal (MD) thalamus, warrants consideration (Barrett & Miller, 2026). The MD thalamus is a critical hub for prefrontal uncertainty processing and representational sharpening (Lam et al., 2025; Mukherjee et al., 2021), and a parallel IT-MD-PFC pathway could theoretically introduce varying latencies. However, a static subcortical bypass struggles to explain the high degree of temporal flexibility observed in our data. Specifically, we found that the PFC-to-IT information flow accelerated by over 50ms specifically when the task was difficult (Figure 10). This demand-dependent ’acceleration’ of the feedback loop—coupled with the qualitative temporal reversal only under uncertainty—is more consistent with an active, top-down intervention than a fixed subcortical bypass. Future studies employing simultaneous thalamic recordings or causal manipulations will be essential to further dissociate the specific contributions of transthalamic versus trans-cortical circuits to the resolution of perceptual ambiguity.

The analysis of information flow as a function of behavioral relevance adds a critical dimension to this interplay. We observed a ’dual-pronged’ strategy: a fast, selective FF mechanism that prioritizes goal-congruent information (Desimone & Duncan, 1995), alongside a stable FB system that maintains the broader cognitive context (Duan et al., 2024). This early prioritization in the feed-forward sweep is consistent with invasive evidence showing that the PFC selectively integrates sensory evidence based on current task rules while effectively gating irrelevant dimensions (Mante et al., 2013; Muhammad et al., 2006). Such a mechanism suggests two distinct functional regimes: *proactive maintenance* and *reactive recruitment*. While the PFC provides a stable ’broad-spectrum’ categorical template regardless of relevance (proactive maintenance), it is temporally prioritized and intensified specifically when sensory signals are too weak for automatic categorization (reactive recruitment). This distinction aligns with modern theories of recurrent vision (Barrett & Miller, 2026) and explains why the PFC-to-IT pathway acts as a constant scaffold that only modulates the hierarchy under conditions of uncertainty.

These results provide a content-resolved bridge to established causal evidence. While prefrontal inactivation is known to impair the recognition of ’challenge’ images (Kar & DiCarlo, 2021), the specific informational currency being exchanged was previously unclear. Our findings suggest this necessity stems from a dynamic reconfiguration: when sensory evidence is ’Close’ to a boundary, the PFC provides the early categorical scaffold required to resolve IT ambiguity, identifying the prefrontal cortex as the essential directional circuit-breaker for perceptual uncertainty (Namima & Pasupathy, 2021; Noroozi et al., 2024; Tang et al., 2018).

An important methodological innovation of the present study lies in our application of Model-Based Representational Connectivity Analysis (RCA). Moving beyond traditional univariate connectivity, which often remains blind to information distributed across neural populations, model-based RCA specifically identifies statistical dependencies based on representational content (Karimi-Rouzbahani et al., 2021, 2022). Critically, whereas conventional methods reveal whether two regions are co-activated, RCA reveals what specific information is being transferred. This allowed us to move from asking whether IT and PFC are connected to precisely quantifying the dynamic flow of multiple, distinct types of information—exemplar details, relational geometries, or abstract categories—dissecting the temporal evolution of bottom-up transmission and top-down modulation. Unlike traditional measures such as Granger Causality, which can be sensitive to trial-to-trial noise and common-input confounds, as they rely on activity rather than information, RCA assesses how the categorical structure of one region informs another. By operating in this ’information space’, the method is less susceptible to session-specific noise and focuses specifically on the transfer of cognitively relevant information content. This robustness makes the framework potentially applicable across diverse modalities or even non-simultaneous datasets, although the simultaneous recordings used here provide the highest level of inferential power for tracking content-specific communication.

Finally, the dynamic reconfiguration of cortical hierarchy, as a function of categorical uncertainty, characterized here offers a critical neurophysiological benchmark for next-generation computational models of vision. While current state-of-the-art architectures like CORnet-S and CORnet-R incorporate recurrence to improve recognition (Kang et al., 2026; Kubilius et al., 2019), they largely rely on fixed temporal loops. Our results suggest that truly human-like flexibility requires a ’heterarchical’ architecture (Pessoa, 2023) that can switch between feedforward and feedback-dominant modes based on internal estimates of uncertainty. This aligns with recent calls for biologically plausible recurrent models that can replicate the demand-dependent shift from feedforward to feedback-driven resolution observed in the primate brain (Barrett & Miller, 2026; Casile et al., 2025; Kang et al., 2026; Tang et al., 2018).

In summary, this study leveraged a novel tripartite analysis of information coding, recurrence, and content-specific information flow to provide a nuanced account of inferotemporal-prefrontal cortical interactions during visual categorization. Our findings not only reaffirm the established hierarchy—with IT rapidly extracting transient, primarily semi-abstract relational features, and PFC maintaining sustained, actively recurrent task-relevant information—but they also unveiled the profound dynamic and content-specific nature of their interplay. We demonstrate that the cortical hierarchy dynamically adapts to cognitive demands. Most significantly, we reveal that under conditions of perceptual uncertainty, the PFC actively initiates a top-down feedback flow to guide sensory processing and resolve ambiguity. By applying state-of-the-art representational connectivity analysis, this work provides a framework for understanding how content-specific, dynamic interactions between sensory and association cortices underlie flexible, resource-efficient cognitive behavior.

## Acknowledgements

HKR was supported by an Australian Research Council DECRA Fellowship DE230100608 and follow-on Newton International Fellowship grants from the UK Royal Society (AL231037 & AL24100035). We thank the Mater Foundation for their support. We would like to thank Professor Earl Miller, Dr Scott Brincat and Dr Jefferson Roy from MIT for providing the monkey dataset, the task and stimulus images and useful feedback on the initial analyses.

## Code and Data Availability

The code for the different analyses, including coding, recurrence and the new model-based Representational Connectivity Analysis (RCA) is available at: https://github.com/HamidKarimi-Rouzbahani/catDog2Bound_Monkey_IT_PFC_Interaction.git. For the data, contact the corresponding authors of the manuscript where the dataset was borrowed from: Roy et al., 2010.

---

**Figure S1.**
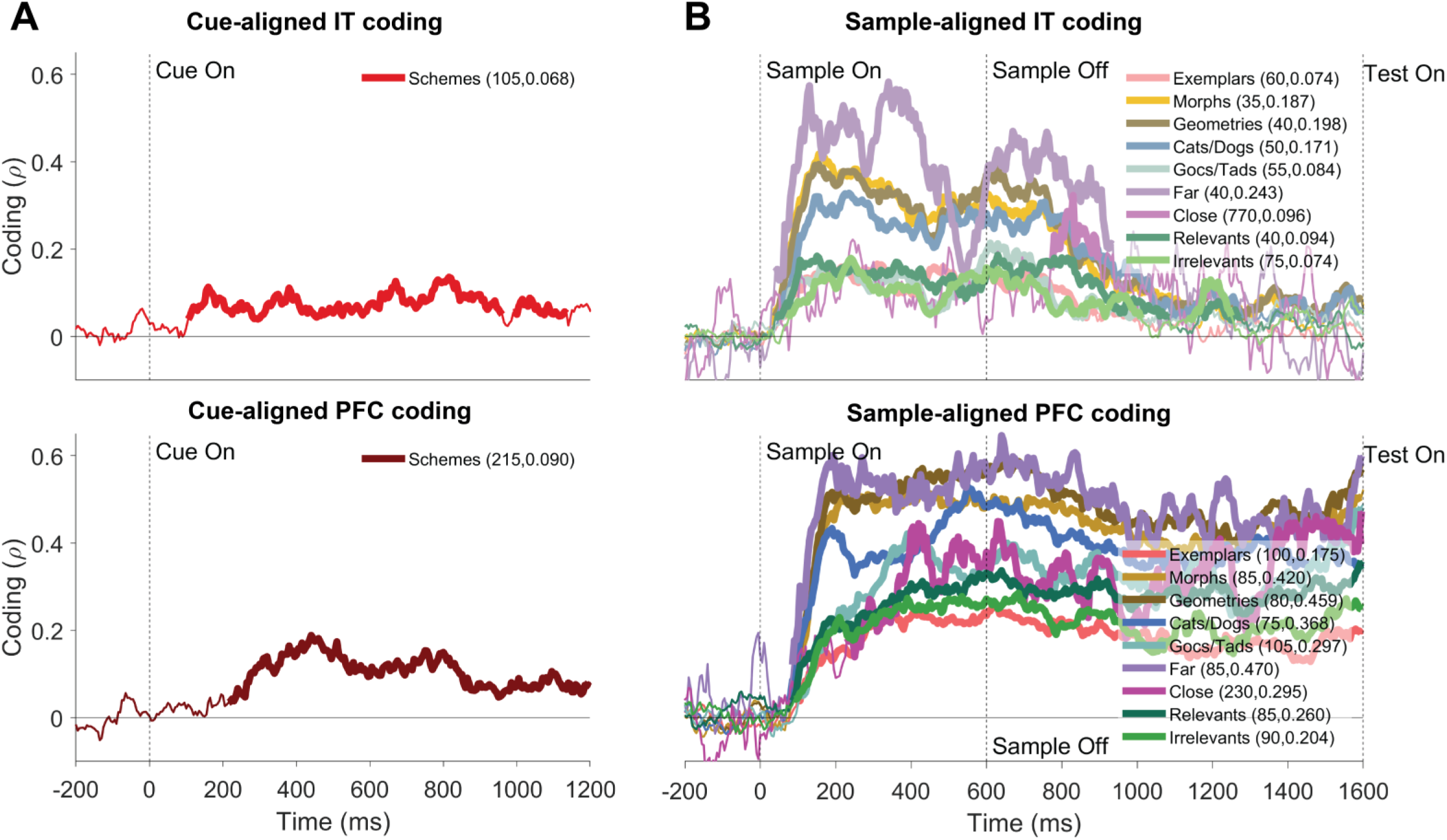
Population-level scheme and stimulus coding. **(A)** Cue-aligned scheme information coding in IT (top) and PFC (bottom). **(B)** Sample-aligned stimulus information coding in IT (top) and PFC (bottom). Different line colors correspond to different conceptual models. The values in paratheses show the onset time of significant neural coding (first number) in milliseconds and the average post-onset area-under-the-curve (AUC, second number). Thickened lines indicate the time points when the coding was significantly above chance (permutation test, p < 0.05; cluster-corrected for FWER). Vertical dashed lines mark critical trial events (i.e., cue, sample onset and offset).

**Figure S2.**
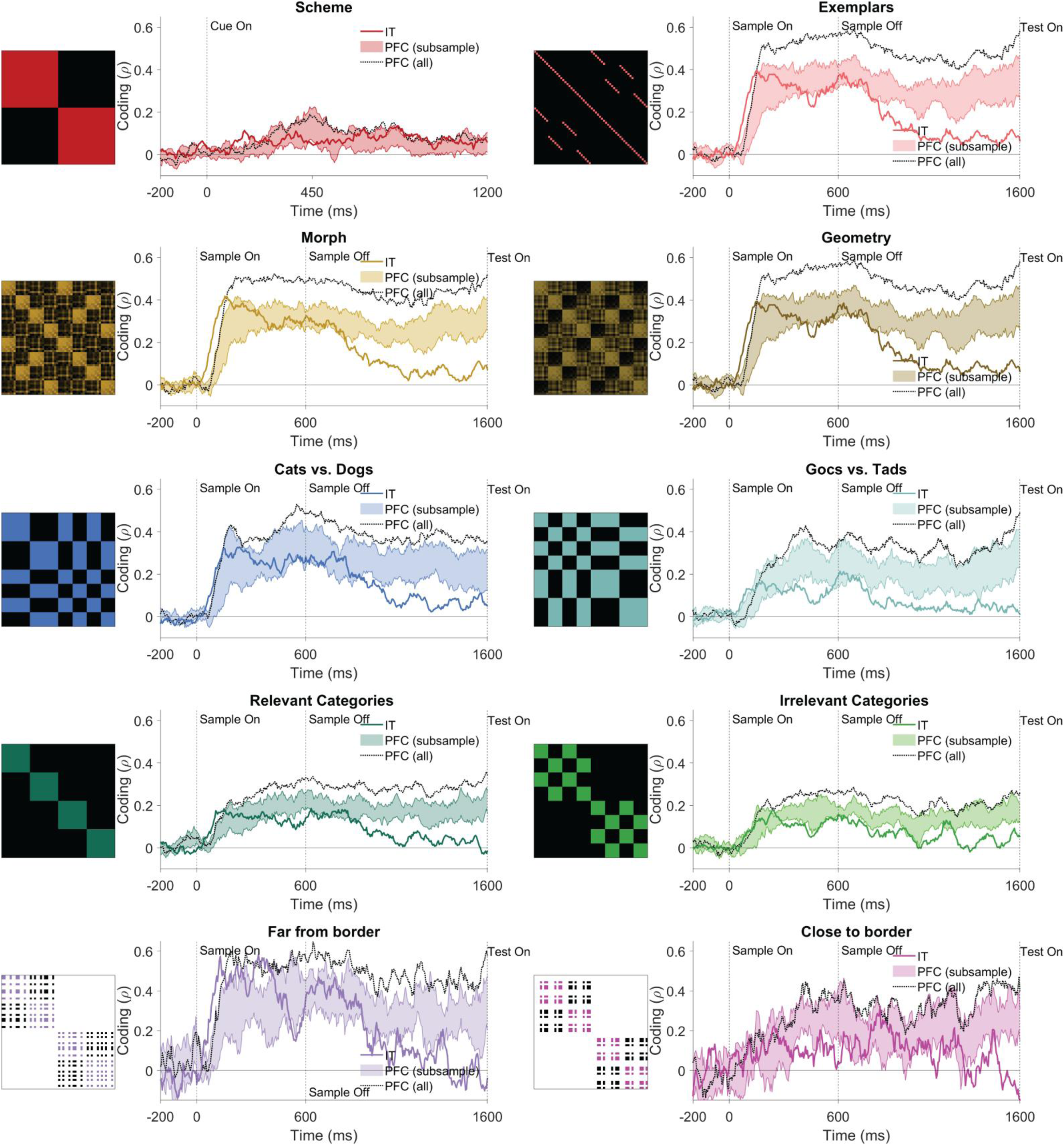
Population-level stimulus coding after subsampling the PFC neurons 10 times. Sample-aligned stimulus information coding in IT (colored), original 549 PFC (black dotted line) and 10 subsampled PFC (colored shading) neurons. Different colors correspond to different conceptual models. PFC neurons were subsampled (from 549 to 154) 10 times to equalize the number of IT neurons and the shadings show the range of coding obtained from the 10 subsampled populations from PFC. Vertical dashed lines mark critical trial events (i.e., cue, sample onset and offset).

**Figure S3.**
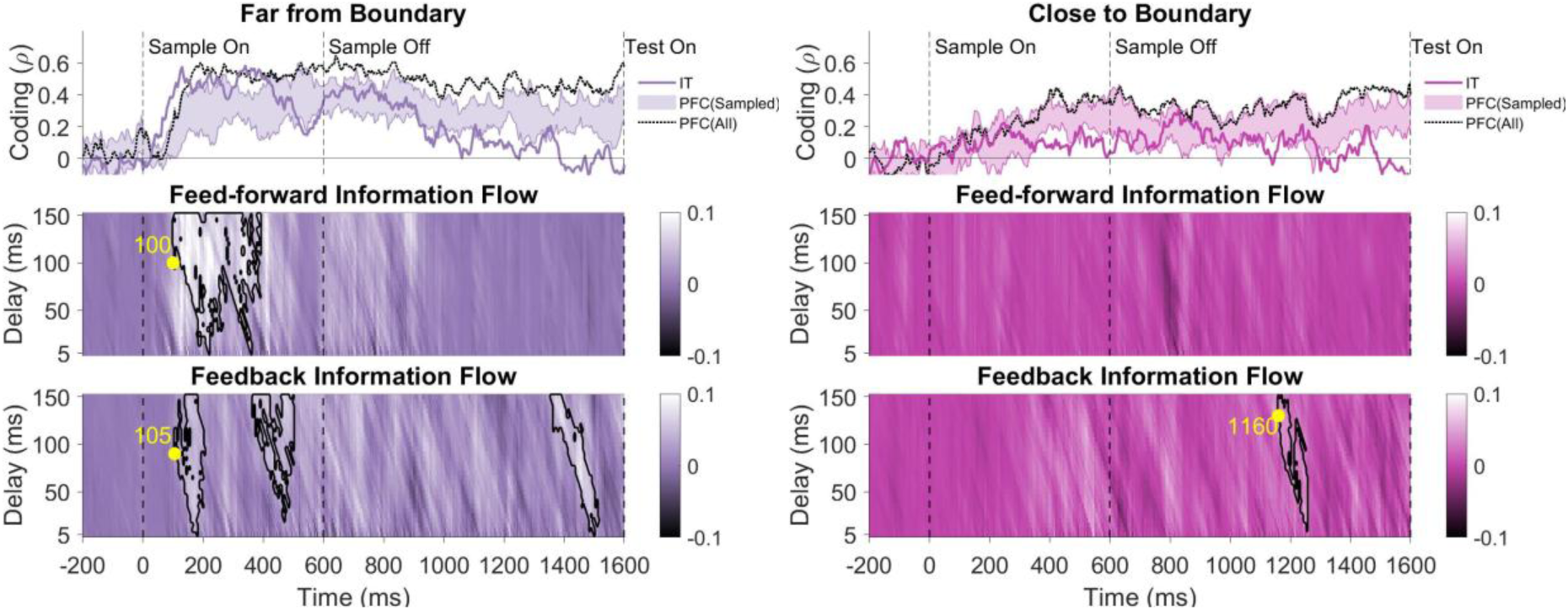
Modulation of sub-sampled information coding and flow with stimulus certainty. Top. (Information Coding) Sample-aligned stimulus information coding in IT (colored), original 549 PFC (black dotted line) and 10 subsampled PFC (shading) neurons (left: Far model; right: Close model). **Middle** (Feed-Forward flow): IT-to-PFC information flow over time (x-axis) and transmission delays (y-axis: 5-150 ms). Black contours enclose significant flows (permutation test, p < 0.05; cluster-corrected for FWER), with yellow dots marking the first significant latencies (values in ms). **Bottom** (Feedback flow): PFC-to-IT information flow, analyzed identically to feed-forward flow, but in the opposite direction. The dots and corresponding numbers show the time point of the first significant coding or flow. Vertical dashed lines refer to response onset.

## References

1. Ahissar, M., & Hochstein, S. (2004). The reverse hierarchy theory of visual perceptual learning. Trends in Cognitive Sciences, 8(10), 457–464. 10.1016/j.tics.2004.08.011

2. Anzellotti, S., & Coutanche, M. N. (2018). Beyond Functional Connectivity: Investigating Networks of Multivariate Representations. Trends in Cognitive Sciences, 22(3), 258–269. 10.1016/j.tics.2017.12.002

3. Ashby, F. G., & Gott, R. E. (1988). Decision rules in the perception and categorization of multidimensional stimuli. J. Exp. Psychol. Learn. Mem. Cogn, 14(1), 33–53,. 10.1037/0278-7393.14.1.33.

4. Ashby, F. G., & Maddox, W. T. (1992). Complex decision rules in categorization: Contrasting novice and experienced performance. J. Exp. Psychol. Hum. Percept. Perform, 18(1), 50–71,. 10.1037/0096-1523.18.1.50.

5. Avarguès-Weber, A., Deisig, N., & Giurfa, M. (2011). Visual cognition in social insects. Annual Review of Entomology, 56, 423–443. 10.1146/annurev-ento-120709-144855

6. Baldassi, C., Alemi-Neissi, A., Pagan, M., DiCarlo, J. J., Zecchina, R., & Zoccolan, D. (2013). Shape Similarity, Better than Semantic Membership, Accounts for the Structure of Visual Object Representations in a Population of Monkey Inferotemporal Neurons. PLOS Comput. Biol, 9(8), 1003167,. 10.1371/journal.pcbi.1003167.

7. Bar, M., Kassam, K. S., Ghuman, A. S., Boshyan, J., Schmid, A. M., Dale, A. M., Hämäläinen, M. S., Marinkovic, K., Schacter, D. L., Rosen, B. R., & Halgren, E. (2006). Top-down facilitation of visual recognition. Proceedings of the National Academy of Sciences, 103(2), 449–454. 10.1073/pnas.0507062103

8. Barrett, L. F., & Miller, E. K. (2026). Categorization is ‘baked’ into the brain. Nature Reviews Neuroscience. 10.1038/s41583-026-01036-2

9. Basti, A., Nili, H., Hauk, O., Marzetti, L., & Henson, R. N. (2020). Multi-dimensional connectivity: A conceptual and mathematical review. NeuroImage, 221, 117179. 10.1016/j.neuroimage.2020.117179

10. Bastos, A. M., & Schoffelen, J.-M. (2016). A Tutorial Review of Functional Connectivity Analysis Methods and Their Interpretational Pitfalls. Frontiers in Systems Neuroscience, 9. 10.3389/fnsys.2015.00175

11. Bastos, A. M., Usrey, W. M., Adams, R. A., Mangun, G. R., Fries, P., & Friston, K. J. (2012). Canonical microcircuits for predictive coding. Neuron, 76(4), 695–711. 10.1016/j.neuron.2012.10.038

12. Brincat, S. L., & Connor, C. E. (2004). Underlying principles of visual shape selectivity in posterior inferotemporal cortex. Nature Neuroscience, 7(8), 880–886. 10.1038/nn1278

13. Buschman, T. J., & Kastner, S. (2015). From behavior to neural dynamics: An integrated theory of attention. Neuron, 88(1), 127–144. 10.1016/j.neuron.2015.09.017

14. Casile, A., Cordier, A., Kim, J. G., Cometa, A., Madsen, J. R., Stone, S., Ben-Yosef, G., Ullman, S., Anderson, W., & Kreiman, G. (2025). Neural correlates of minimal recognizable configurations in the human brain. Cell Reports, 44(3), 115429. 10.1016/j.celrep.2025.115429

15. Cole, M. W., Ito, T., & Braver, T. S. (2016). The Behavioral Relevance of Task Information in Human Prefrontal Cortex. Cerebral Cortex (New York, NY), 26(6), 2497–2505. 10.1093/cercor/bhv072

16. Debes, S. R., & Dragoi, V. (2023). Suppressing feedback signals to visual cortex abolishes attentional modulation. Science. (world). 10.1126/science.ade1855

17. Desimone, R. (1996). Neural mechanisms for visual memory and their role in attention. Proceedings of the National Academy of Sciences of the United States of America, 93(24), 13494–13499. 10.1073/pnas.93.24.13494

18. Desimone, R., Albright, T., Gross, C., & Bruce, C. (1984). Stimulus-selective properties of inferior temporal neurons in the macaque. The Journal of Neuroscience, 4(8), 2051–2062. 10.1523/JNEUROSCI.04-08-02051.1984

19. Desimone, R., & Duncan, J. (1995). Neural mechanisms of selective visual attention. Annual Review of Neuroscience, 18, 193–222. 10.1146/annurev.ne.18.030195.001205

20. DiCarlo, J. J., Zoccolan, D., & Rust, N. C. (2012). How does the brain solve visual object recognition? Neuron, 73(3), 415–434. 10.1016/j.neuron.2012.01.010

21. Dienes, Z. (2014). Using Bayes to get the most out of non-significant results. Frontiers in Psychology, 5. 10.3389/fpsyg.2014.00781

22. Dijkstra, N., Zeidman, P., Ondobaka, S., van Gerven, M. a. J., & Friston, K. (2017). Distinct Top-down and Bottom-up Brain Connectivity During Visual Perception and Imagery. Scientific Reports, 7(1), 5677. 10.1038/s41598-017-05888-8

23. Duan, Y., Zhan, J., Gross, J., Ince, R. A. A., & Schyns, P. G. (2024). Pre-frontal cortex guides dimension-reducing transformations in the occipito-ventral pathway for categorization behaviors. Current Biology, 34(15), 3392–3404.e5. 10.1016/j.cub.2024.06.050

24. Duncan, J. (2001). An adaptive coding model of neural function in prefrontal cortex. Nature Reviews Neuroscience, 2(11), 820–829. 10.1038/35097575

25. Felleman, D. J., & Van Essen, D. C. (1991). Distributed Hierarchical Processing in the Primate Cerebral Cortex. Cerebral Cortex, 1(1), 1–47. 10.1093/cercor/1.1.1-a

26. Freedman, D. J., Riesenhuber, M., Poggio, T., & Miller, E. K. (2001). Categorical representation of visual stimuli in the primate prefrontal cortex. *Science (New York*, N.Y.), 291(5502), 312–316. 10.1126/science.291.5502.312

27. Freedman, D. J., Riesenhuber, M., Poggio, T., & Miller, E. K. (2003). A Comparison of Primate Prefrontal and Inferior Temporal Cortices during Visual Categorization. Journal of Neuroscience, 23(12), 5235–5246. 10.1523/JNEUROSCI.23-12-05235.2003

28. Friston, K. (2005). A theory of cortical responses. Philosophical Transactions of the Royal Society B: Biological Sciences, 360(1456), 815–836. 10.1098/rstb.2005.1622

29. Funahashi, S., Bruce, C. J., & Goldman-Rakic, P. S. (1989). Mnemonic coding of visual space in the monkey’s dorsolateral prefrontal cortex. Journal of Neurophysiology, 61(2), 331–349. 10.1152/jn.1989.61.2.331

30. Fuster, J. M. (2001). The Prefrontal Cortex—An Update: Time Is of the Essence. Neuron, 30(2), 319–333. 10.1016/S0896-6273(01)00285-9

31. Fuster, J. M., & Alexander, G. E. (1971). Neuron activity related to short-term memory. Science (New York, N.Y.), 173(3997), 652–654. 10.1126/science.173.3997.652

32. Fyall, A. M., El-Shamayleh, Y., Choi, H., Shea-Brown, E., & Pasupathy, A. (2017). Dynamic representation of partially occluded objects in primate prefrontal and visual cortex. eLife, 6, e25784. 10.7554/eLife.25784

33. Gilbert, C. D., & Li, W. (2013). Top-down influences on visual processing. Nature Reviews Neuroscience, 14(5), 350–363. 10.1038/nrn3476

34. Goddard, E., Carlson, T. A., Dermody, N., & Woolgar, A. (2016). Representational dynamics of object recognition: Feedforward and feedback information flows. NeuroImage, 128, 385–397,. 10.1016/j.neuroimage.2016.01.006.

35. Goltstein, P. M., Reinert, S., Bonhoeffer, T., & Hübener, M. (2021). Mouse visual cortex areas represent perceptual and semantic features of learned visual categories. Nature Neuroscience, 24(10), 1441–1451. 10.1038/s41593-021-00914-5

36. Gregoriou, G. G., Gotts, S. J., Zhou, H., & Desimone, R. (2009). High-frequency, long-range coupling between prefrontal and visual cortex during attention. *Science (New York*, N.Y.), 324(5931), 1207–1210. 10.1126/science.1171402

37. Gross, C. G. (2002). Genealogy of the ‘Grandmother Cell,.’ The Neuroscientist, 8(5), 512–518. 10.1177/107385802237175.

38. Hebart, M. N., Bankson, B. B., Harel, A., Baker, C. I., & Cichy, R. M. (2018). The representational dynamics of task and object processing in humans. eLife, 7, e32816. 10.7554/eLife.32816

39. Herrnstein, R. J., & Loveland, D. H. (1964). COMPLEX VISUAL CONCEPT IN THE PIGEON. *Science (New York*, N.Y.), 146(3643), 549–551. 10.1126/science.146.3643.549

40. Jackson, J. B., Feredoes, E., Rich, A. N., Lindner, M., & Woolgar, A. (2021). Concurrent neuroimaging and neurostimulation reveals a causal role for dlPFC in coding of task-relevant information. Communications Biology, 4(1), 588. 10.1038/s42003-021-02109-x

41. Jackson, J., Rich, A. N., Williams, M. A., & Woolgar, A. (2017). Feature-selective Attention in Frontoparietal Cortex: Multivoxel Codes Adjust to Prioritize Task-relevant Information. Journal of Cognitive Neuroscience, 29(2), 310–321. 10.1162/jocn_a_01039

42. Jeffreys, H. (1998). The theory of probability (3rd ed.). Oxford University Press.

43. Kang, B., Midler, B., Chen, F., & Druckmann, S. (2026). Recurrent connections facilitate occluded object recognition by explaining-away. Nature Communications, 17(1), 2225. 10.1038/s41467-026-68806-5

44. Kar, K., & DiCarlo, J. J. (2021). Fast Recurrent Processing via Ventrolateral Prefrontal Cortex Is Needed by the Primate Ventral Stream for Robust Core Visual Object Recognition. Neuron, 109(1), 164–176.e5. 10.1016/j.neuron.2020.09.035

45. Kar, K., Kubilius, J., Schmidt, K., Issa, E. B., & DiCarlo, J. J. (2019). Evidence that recurrent circuits are critical to the ventral stream’s execution of core object recognition behavior. Nature Neuroscience, 22(6), 974–983. 10.1038/s41593-019-0392-5

46. Karimi-Rouzbahani, H., Ramezani, F., Woolgar, A., Rich, A., & Ghodrati, M. (2021). Perceptual difficulty modulates the direction of information flow in familiar face recognition. NeuroImage, 233, 117896. 10.1016/j.neuroimage.2021.117896

47. Karimi-Rouzbahani, H., Woolgar, A., Henson, R., & Nili, H. (2022). Caveats and Nuances of Model-Based and Model-Free Representational Connectivity Analysis. Frontiers in Neuroscience, 16. 10.3389/fnins.2022.755988

48. Kiani, R., Esteky, H., Mirpour, K., & Tanaka, K. (2007). Object Category Structure in Response Patterns of Neuronal Population in Monkey Inferior Temporal Cortex. Journal of Neurophysiology, 97(6), 4296–4309. 10.1152/jn.00024.2007

49. King, J.-R., & Dehaene, S. (2014). Characterizing the dynamics of mental representations: The temporal generalization method. Trends in Cognitive Sciences, 18(4), 203–210. 10.1016/j.tics.2014.01.002

50. Kreiman, G., & Serre, T. (2020). Beyond the feedforward sweep: Feedback computations in the visual cortex. Annals of the New York Academy of Sciences, 1464(1), 222–241. 10.1111/nyas.14320

51. Kriegeskorte, N., Mur, M., & Bandettini, P. A. (2008). Representational similarity analysis—Connecting the branches of systems neuroscience. Frontiers in Systems Neuroscience, 2. 10.3389/neuro.06.004.2008

52. Kubilius, J., Schrimpf, M., Kar, K., Hong, H., Majaj, N. J., Rajalingham, R., Issa, E. B., Bashivan, P., Prescott-Roy, J., Schmidt, K., Nayebi, A., Bear, D., Yamins, D. L. K., & DiCarlo, J. J. (2019). *Brain-Like Object Recognition with High-Performing Shallow Recurrent ANNs* (Version 2). arXiv. 10.48550/ARXIV.1909.06161

53. Lam, N. H., Mukherjee, A., Wimmer, R. D., Nassar, M. R., Chen, Z. S., & Halassa, M. M. (2025). Prefrontal transthalamic uncertainty processing drives flexible switching. Nature, 637(8044), 127–136. 10.1038/s41586-024-08180-8

54. Lee, M. D., & Wagenmakers, E.-J. (2005). Bayesian statistical inference in psychology: Comment on Trafimow (2003). Psychological Review, 112(3), 662–668. 10.1037/0033-295X.112.3.662

55. Lee, S.-H., & Baker, C. I. (2016). Multi-Voxel Decoding and the Topography of Maintained Information During Visual Working Memory. Frontiers in Systems Neuroscience, 10. 10.3389/fnsys.2016.00002

56. Liu, Z.-Q., Luppi, A. I., Hansen, J. Y., Tian, Y. E., Zalesky, A., Yeo, B. T. T., Fulcher, B. D., & Misic, B. (2025). Benchmarking methods for mapping functional connectivity in the brain. Nature Methods, 22(7), 1593–1602. 10.1038/s41592-025-02704-4

57. Majaj, N. J., Hong, H., Solomon, E. A., & DiCarlo, J. J. (2015). Simple Learned Weighted Sums of Inferior Temporal Neuronal Firing Rates Accurately Predict Human Core Object Recognition Performance. The Journal of Neuroscience, 35(39), 13402–13418. 10.1523/JNEUROSCI.5181-14.2015

58. Mansouri, F. A., Tanaka, K., & Buckley, M. J. (2009). Conflict-induced behavioural adjustment: A clue to the executive functions of the prefrontal cortex. Nature Reviews. Neuroscience, 10(2), 141–152. 10.1038/nrn2538

59. Mante, V., Sussillo, D., Shenoy, K. V., & Newsome, W. T. (2013). Context-dependent computation by recurrent dynamics in prefrontal cortex. Nature, 503(7474), 78–84. 10.1038/nature12742

60. Meyers, E. M., Freedman, D. J., Kreiman, G., Miller, E. K., & Poggio, T. (2008). Dynamic Population Coding of Category Information in Inferior Temporal and Prefrontal Cortex. Journal of Neurophysiology, 100(3), 1407–1419. 10.1152/jn.90248.2008

61. Miller, E. K., & Cohen, J. D. (2001). An integrative theory of prefrontal cortex function. Annual Review of Neuroscience, 24, 167–202. 10.1146/annurev.neuro.24.1.167

62. Miller, E. K., Lundqvist, M., & Bastos, A. M. (2018). Working Memory 2.0. Neuron, 100(2), 463–475. 10.1016/j.neuron.2018.09.023

63. Minda, J. P., Roark, C. L., Kalra, P., & Cruz, A. (2024). Single and multiple systems in categorization and category learning. Nat. Rev. Psychol, 3(8), 536–551,. 10.1038/s44159-024-00336-7.

64. Moore, T., & Armstrong, K. M. (2003). Selective gating of visual signals by microstimulation of frontal cortex. Nature, 421(6921), 370–373. 10.1038/nature01341

65. Muhammad, R., Wallis, J. D., & Miller, E. K. (2006). A comparison of abstract rules in the prefrontal cortex, premotor cortex, inferior temporal cortex, and striatum. J. Cogn. Neurosci, 18(6), 974–989,. 10.1162/jocn.2006.18.6.974.

66. Mukherjee, A., Lam, N. H., Wimmer, R. D., & Halassa, M. M. (2021). Thalamic circuits for independent control of prefrontal signal and noise. Nature, 600(7887), 100–104. 10.1038/s41586-021-04056-3

67. Namima, T., & Pasupathy, A. (2021). Encoding of Partially Occluded and Occluding Objects in Primate Inferior Temporal Cortex. The Journal of Neuroscience, 41(26), 5652–5666. 10.1523/JNEUROSCI.2992-20.2021

68. Nili, H., Wingfield, C., Walther, A., Su, L., Marslen-Wilson, W., & Kriegeskorte, N. (2014). A Toolbox for Representational Similarity Analysis. PLoS Computational Biology, 10(4), e1003553. 10.1371/journal.pcbi.1003553

69. Noroozi, J., Rezayat, E., & Dehaqani, M.-R. A. (2024). Frontotemporal network contribution to occluded face processing. Proceedings of the National Academy of Sciences, 121(48), e2407457121. (world). 10.1073/pnas.2407457121

70. Pessoa, L. (2023). The Entangled Brain. J. Cogn. Neurosci, 35(3), 349–360,. 10.1162/jocn_a_01908.

71. Rajaei, K., Mohsenzadeh, Y., Ebrahimpour, R., & Khaligh-Razavi, S.-M. (2019). Beyond core object recognition: Recurrent processes account for object recognition under occlusion. PLOS Computational Biology, 15(5), e1007001. 10.1371/journal.pcbi.1007001

72. Reinert, S., Hübener, M., Bonhoeffer, T., & Goltstein, P. M. (2021). Mouse prefrontal cortex represents learned rules for categorization. Nature, 593(7859), 411–417. 10.1038/s41586-021-03452-z

73. Riesenhuber, M., & Poggio, T. (1999). Hierarchical models of object recognition in cortex. Nature Neuroscience, 2(11), 1019–1025. 10.1038/14819

74. Riesenhuber, M., & Poggio, T. (2000). Models of object recognition. Nature Neuroscience, 3(11), 1199–1204. 10.1038/81479

75. Roberts, W. A., & Mazmanian, D. S. (1988). Concept learning at different levels of abstraction by pigeons, monkeys, and people. Journal of Experimental Psychology: Animal Behavior Processes, 14(3), 247–260. 10.1037/0097-7403.14.3.247

76. Rouder, J. N., Morey, R. D., Speckman, P. L., & Province, J. M. (2012). Default Bayes factors for ANOVA designs. Journal of Mathematical Psychology, 56(5), 356–374. 10.1016/j.jmp.2012.08.001

77. Roy, J. E., Riesenhuber, M., Poggio, T., & Miller, E. K. (2010). Prefrontal Cortex Activity during Flexible Categorization. The Journal of Neuroscience, 30(25), 8519–8528. 10.1523/JNEUROSCI.4837-09.2010

78. Rust, N. C., & DiCarlo, J. J. (2010). Selectivity and Tolerance (“Invariance”) Both Increase as Visual Information Propagates from Cortical Area V4 to IT. Journal of Neuroscience, 30(39), 12978–12995. 10.1523/JNEUROSCI.0179-10.2010

79. Seger, C. A., & Miller, E. K. (2010). Category Learning in the Brain. Annual Review of Neuroscience, 33, 203–219. 10.1146/annurev.neuro.051508.135546

80. Shelton, C. R. (2000). Morphable Surface Models. International Journal of Computer Vision, 38(1), 75–91. 10.1023/A:1008170818506

81. Tanaka, K. (1996). Inferotemporal cortex and object vision. Annual Review of Neuroscience, 19, 109–139. 10.1146/annurev.ne.19.030196.000545

82. Tang, H., Schrimpf, M., Lotter, W., Moerman, C., Paredes, A., Ortega Caro, J., Hardesty, W., Cox, D., & Kreiman, G. (2018). Recurrent computations for visual pattern completion. Proceedings of the National Academy of Sciences, 115(35), 8835–8840. 10.1073/pnas.1719397115

83. Tomita, H., Ohbayashi, M., Nakahara, K., Hasegawa, I., & Miyashita, Y. (1999). Top-down signal from prefrontal cortex in executive control of memory retrieval. Nature, 401(6754), 699–703. 10.1038/44372

84. Vogels, R. (1999). Categorization of complex visual images by rhesus monkeys. Part 2: Single-cell study. The European Journal of Neuroscience, 11(4), 1239–1255. 10.1046/j.1460-9568.1999.00531.x

85. Vogels, R. (2022). More Than the Face: Representations of Bodies in the Inferior Temporal Cortex. Annual Review of Vision Science, 8(1), 383–405. 10.1146/annurev-vision-100720-113429

86. Wallis, J. D., Anderson, K. C., & Miller, E. K. (2001). Single neurons in prefrontal cortex encode abstract rules. Nature, 411(6840), 953–956. 10.1038/35082081

87. Wasserman, E. A., Kiedinger, R. E., & Bhatt, R. S. (1988). Conceptual behavior in pigeons: Categories, subcategories, and pseudocategories. Journal of Experimental Psychology: Animal Behavior Processes, 14(3), 235–246. 10.1037/0097-7403.14.3.235

88. White, I. M., & Wise, S. P. (1999). Rule-dependent neuronal activity in the prefrontal cortex. Experimental Brain Research, 126(3), 315–335. 10.1007/s002210050740

89. Wyttenbach, R. A., May, M. L., & Hoy, R. R. (1996). Categorical perception of sound frequency by crickets. *Science (New York*, N.Y.), 273(5281), 1542–1544. 10.1126/science.273.5281.1542

90. Zellner, A., & Siow, A. (1980). Posterior odds ratios for selected regression hypotheses. Trabajos de Estadistica Y de Investigacion Operativa, 31(1), 585–603. 10.1007/BF02888369

